# A Curvature-Enhanced Random Walker Segmentation Method for Detailed Capture of 3D Cell Surface Membranes

**DOI:** 10.1101/720177

**Authors:** E. Josiah Lutton, Sharon Collier, Till Bretschneider

## Abstract

High-resolution 3D microscopy is a fast advancing field and requires new techniques in image analysis to handle these new datasets. In this work, we focus on detailed 3D segmentation of *Dictyostelium* cells undergoing macropinocytosis captured on an iSPIM microscope. We propose a novel random walker-based method with a curvature-based enhancement term, with the aim of capturing fine protrusions, such as filopodia and deep invaginations, such as macropinocytotic cups, on the cell surface. We tested our method on both real and synthetic 3D image volumes, demonstrating that the inclusion of the curvature enhancement term can improve the segmentation of the aforementioned features. We show that our method performs better than other state of the art segmentation methods in 3D images of *Dictyostelium* cells, and performs competitively against CNN-based methods in challenge datasets, demonstrating the ability to obtain accurate segmentations without the requirement of large training datasets. We also present an automated seeding method for microscopy data, which, combined with the curvature-enhanced random walker method, enables the segmentation of large time series with minimal input from the experimenter.

## I. Introduction

RECENT advances in 3D microscopy have enabled the detailed capture of large time series of cells, posing challenges of accurately segmenting them in high resolution. In this work we focus on the segmentation of *Dictyostelium* cells showing complex cell deformations during vesicular uptake of nutrients from extracellular fluid, a process called macropinocytosis. Macropinocytosis is an important component of cancer cell feeding [1] and antigen processing of macrophages [2]. It involves the formation of highly concave invaginations of the cell membrane, macropinocytotic cups, that are subsequently shaped into vesicles. Filopodia, fine protrusions of the cell membrane with highly convex tips, have recently been assigned a role in macrophage macropinocytosis, too [3]. In order to better understand the role of both of these structures in macropinocytosis, accurate segmentations are required [4], [5], which is difficult due to their high curvature. The cells examined expressed LifeAct-GFP, a commonly used fluorescent F-actin marker present in both structures.

Fast 3D light sheet imaging has recently become the method of choice to capture the fast dynamics of macropinocytotic cups and filopodia [3], [6]. Deconvolution of the 3D light sheet data is employed to sharpen images, but theoretically optimum deblurring is hardly ever achieved in practice. Remaining blur specifically compromises highly curved structures, causing protrusions to be truncated or lost during segmentation, while invaginations tend to lose depth. We extend the random walker segmentation, a standard method for segmenting objects in 2- and 3-dimensional images [7], [8], by incorporating a curvature enhancement term to recover these structures. Additionally, we present a method for automated seed selection for the random walker, which utilizes the fluorescence pattern of the F-Actin marker. These methods were tested on both real and synthetic image volumes, showing that the addition of curvature enhancement to the random walker method can provide a marked improvement in the segmentation of detailed structures such as those mentioned above. We compared our method with four state of the art segmentation methods: a pretrained convolutional neural network [9], the random forest pixel classifier [10], the power watershed [11], and band pass segmentation [12], and show that our method outperforms all four in real cell images. Additionally, our methods were externally evaluated by the Cell Tracking Challenge (https://www.celltrackingchallenge.net) [12], [13], demonstrating that our method performs competitively against deep learning-based techniques.

## II. Related Work

One of the simplest approaches to image segmentation is to apply a threshold to the image, either by using a globally [14], or locally [15] defined threshold. Unfortunately, fluorescent markers in cells do not tend to produce uniformly high fluorescence throughout the cell, and many only provide partial coverage for the cell membrane, including F-actin markers, which are studied here. This means that simple thresholding is generally insufficient. An option for improving the result of thresholding is to use pre- and post-processing steps. One promising method of this kind applies intensity clipping and a band pass filter prior to thresholding, and then applies a morphological fill operator to the thresholded image [12].

Segmentation methods using machine learning and in particular convolutional neural networks have recently gained popularity [16]–[18]. One issue with implementing these methods is that they rely on a large set of manually annotated training data, which is generally impractical to obtain for 3D images. One method employed to overcome this limitation is to use established segmentation methods to generate the training annotations [19]. This still requires manual verification by the user, however, and is still dependent on the original segmentation method. Another method is to utilize synthetic data to increase the amount of training data [9]. While some data can be generated synthetically through model-based [20], [21] or deep learning-based [22], [23] methods, there is insufficient data of this type available for the cells analyzed here. Machine learning methods that employ pixel classifiers, such as the random forest pixel classifier [10] are more suitable for our data, since they can be trained on a single partially annotated image stack.

A popular method for segmenting whole cells that is related to the random walker is the use of active meshes [24]–[26], which may also be computed implicitly as level sets [27]. This method aims to find the surface that optimizes an energy function dependent on image intensity inside and outside of the surface, and on the geometry of the surface, and is related to the random walker as this energy minimization can be formulated in terms of the graph cuts method mentioned below [28].

The random walker is a commonly-used method of supervised image segmentation [7], [8]. This method is part of a broader family [11], [29] of graph-based segmentation methods, which includes graph cuts [30] and watersheds [31]. These methods model the image as a graph *G* = (*V, E*) where the vertices *V* = {*v_i_*} corresponding to image voxels (or pixels for 2D images) and edges *E* = {*e_ij_*} corresponding to the adjacency relationship of the voxels, with *e_ij_* representing the edge between *v_i_* and *v_j_*. The main idea is to take an input set of voxels marked as foreground and background, referred to as seeds, and expand these sets based on the weighted graph to classify all voxels as either foreground or background. For a given edge weighting *W* = {*w_ij_*}, the aim of a graph-based segmentation is to find a function *x* on *G* that minimizes an energy term of the form [11]

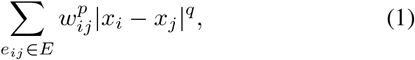

where *x_i_* = *x*(*v_i_*), *x_i_* = 1 at foreground seeds, and *x_i_* = 0 at background seeds. The segmented foreground is given by points with *x >* 0.5. The choice of *q* and *p* determine which segmentation method is being used: *q* = *p* = 1 corresponds to graph cuts [29]; *q* = *p* = 2 corresponds to the random walker [29]; *q* = *p* → *∞* corresponds to shortest paths [29]; *q* = 1*, p* → *∞* corresponds to the watershed [11], [32]; and *q* > 1, *p* → *∞* corresponds to the recently-developed power watershed [11]. Extensions to these methods include adding prior information to the energy function [8], [33], the addition of auxiliary nodes [34], [35], and modification of the edge weighting *w_ij_* [36], [37], which is typically based on image gradients.

Curvature was used previously by M’hiri *et al.* [36] in the weighting of the random walker in order to enhance segmentation of blood vessels in 3D images. This method was facilitated by using a measure of “vesselness” [38] in the input image volume to inform the weighting of the system. This method is not readily applicable to segmenting individual cells, since the structures of interest do not necessarily conform to an easily-defined shape. Another use of curvature in vessel segmentation is in a regularization term for the fast marching segmentation algorithm [39], with the aim of avoiding high-curvature surfaces.

Our methods utilize a model of the random walker based on the discretisation of a non-linear diffusion system [7], [40]. This model allows the addition of a mean curvature term to the model equation. Applications of mean curvature flow in image analysis include image enhancement [41], [42] and active contour-based segmentation methods [43]–[45]. An example that is closely related to the present work is a method of image enhancement proposed by Malladi and Sethian [42], which uses a function of curvature that takes on positive or negative values of curvature depending on the local image properties. This function is similar in construction to the curvature term in Eq. 7 below. Related to mean curvature flow is Willmore flow [46], which has also been used in level set segmentation methods [47]. In 2D systems the Willmore flow is equivalent to minimization of Euler’s elastica energy [48], which has previously been employed in the weighting of graph cut segmentation [49], [50]. These curvature-based segmentation methods use curvature to stabilize the boundaries of segmentation, which is the reverse of the effect produced by the methods presented here; we employ a term that effectively reverses the flow of curvature to improve the segmentation of protrusions and invaginations.

The random walker is highly dependent on the initial seeding [49]. Previous implementations of the random walker algorithm have used manual [7], semi-automated [8], and fully-automated seed selection [51]. Here we present an automated method of seed selection based on Phansalkar thresholding [15].

## III. Materials and Methods

### A. The curvature-enhanced random walker method

Random walker segmentation can be modeled as the steady state of a discretisation of the non-linear diffusion system [7], [40]:

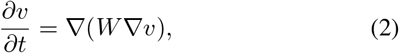

with *∂v/∂n* = 0 at the volume boundary with normal *n*, and subject to the constraints

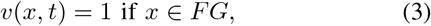

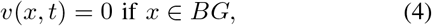

where *FG* and *BG* are the sets of foreground and background seed voxels, respectively, and *W* is the diffusion weighting function, defined discretely between two voxels *x* and *y*, as

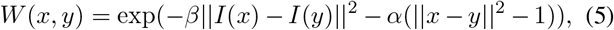

where ||·|| is the Euclidean norm, *I*(*x*) and *I*(*y*) are the input image intensities at *x* and *y* respectively, and *β* and *α* are parameters to be fixed. We set *α* = *β /*255^2^ in accordance with previous work by Du *et al.* [8], with the scale factor of 1*/*255^2^ accounting for scaling from 8-bit image stacks used by Du *et al.* [8] to the range [0, 1], which is used here. Preliminary testing showed that a wide range of values for *α* could be used with similar results. Weights are computed for 18-connected neighborhoods. The discretised form of Eq. 2 for the point *x* is

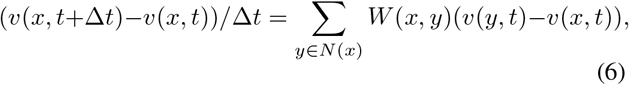

Where *N* (*x*) is the 18-connected neighborhood of *x*, and time step Δ*t* < max(*W*)*/*18 to satisfy the Courant-Friedrichs-Lewy (CFL) condition for numerical stability [52]. The equilibrium values of *v* are computed using the forward Euler method, with the segmented foreground (in the absence of curvature enhancement) given by voxels with *v >* 0.5 [40].

The curvature-enhanced random walker is defined by the system

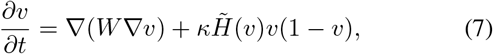

subject to constraints 3 and 4, where *κ* is a fixed parameter, *W* is as defined in Eq. 5, and

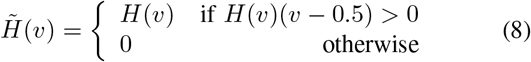

where

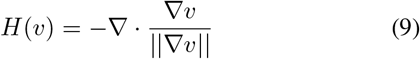

is the mean curvature. The time step is taken to be the same as in the random walker, since, for the values of *κ* studied here, the numerical error introduced by the curvature term is an order of magnitude smaller than that produced by the diffusive term, and therefore stability holds given a time step sufficiently below the maximum required to satisfy the CFL condition for diffusion. This assumption was tested by comparing the results with those generated using smaller time steps, which yielded the same result, as expected. As with the standard random walker, equilibrium values of *v* are calculated and the segmented foreground is given by voxels with *v* > 0.5.

The gradients in Eq. 9 are approximated in each direction using an extension of the 3D Sobel filter, which is given in the *x*-direction as a smoothing in *y* and *z* by applying the 1D filter (1, 4, 6, 4, 1)*/*16 in both *y*- and *z*-directions, Followed by a differencing filter in *x* with radius 3, given by (−1, 0, 0, 0, 0, 0, 1)*/*6. The formulation for *y*- and *z*-directions are defined similarly. The expanded smoothing radius is required because the divergence of the normal directions is highly sensitive to noise. The differences are taken at a distance of 3 voxels to reduce sensitivity to small fluctuations in the shape of the isosurface. Note that this definition assumes isotropic resolution in all directions, which may not be the case for microscopy image stacks. See Section III-D for details on the handling of anisotropy in each of the datasets studied. The number of operations involved in this computation are much higher than for the finite differences in the random walker, and therefore increase computation time. In our implementation we were able to improve the speed by only computing the curvature every 10 time steps, which had a negligible impact on the resulting segmentation.

The curvature-enhanced random walker segmentation is implemented on a GPU as follows. Initially, the equilibrium values *v*_1_ of the standard random walker system are computed, with initial conditions *v*(*x,* 0) = 0.5 for all *x* not in either of the seed sets. The equilibrium values *v*_2_ of the curvature-enhanced diffusion system are subsequently computed with initial conditions *v*(*x,* 0) = *v*_1_. The segmented object is given by the set of voxels with *v*_2_ *>* 0.5. Note that, for large values of *κ*, this system becomes unstable and equilibrium is not reached. In these cases, simulations are terminated after a fixed number of time steps. Source code for this method can be found at https://pilip.lnx.warwick.ac.uk/TMI_2020/.

### B. Seed selection

Automated seed selection is performed on microscopy data based on the Phansalkar threshold [15]. We outline the thresholding method here, and a full description can be found in the supplementary material. Note that, in the following, images are assumed to have isotropic resolution. See Section III-D for details on how anisotropic resolution is handled in each dataset. The Phansalkar threshold assigns a threshold value to each voxel based on the mean and standard deviation of the intensities in a local neighborhood. We adapt the Phansalkar threshold parameter values to each image stack by estimating the local mean and standard deviation for foreground and background voxels. The aim of this adaptive method is to minimize the threshold for foreground voxels, making them more likely to be selected as foreground, and, conversely, maximize the threshold for background voxels.

Background seeds are obtained by applying a spherical dilation operator of radius 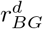 voxels to the binary volume obtained by the Phansalkar thresholding, filling holes, applying a spherical erosion operator of radius 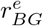 and inverting the resulting binary image volume. An example of the result of this process is shown in Fig. 2C.

Two methods are used for selecting foreground seeds for the data presented in Section III-D. The first is the same method as for the background seed selection, without the final inversion, with dilation and erosion radii being 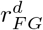 and 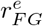 respectively. In the second method, foreground seeds are selected to represent maximal surfaces in the image volume. Gradients are computed using a Sobel operator (Fig. 2D). Each voxel is assigned the direction of the largest magnitude gradient within a 26-connected neighborhood for the purposes of comparison (Fig 2E). Magnitudes along the assigned direction are linearly interpolated from the neighboring voxels, and a voxel is labeled as a turning point if its gradient magnitude is lower than these interpolated magnitudes. Turning points with a negative Laplacian are labeled maximal. This set of maximal points is restricted to points marked as foreground by the Phansalkar thresholding algorithm. To further improve the robustness of this algorithm, the maximal points are grouped into (6-)connected components. The largest connected component is labeled as foreground. Any other connected components larger than one voxel with mean intensity greater than the mean intensity of the largest connected component are also labeled as foreground. An example of the result of this process is shown in Fig. 2C. Parameter values for seed selection in all datasets are available in the supplementary material.

As a result of this seeding method, the edges of the image volume tend to be populated with background seeds, meaning that the volume to be segmented is generally smaller than the input image volume. Accordingly, we automatically crop all images to the smallest bounding cube with background seeds populating the edge planes, in order to reduce processing times.

### C. Comparison with other segmentation methods

We compared the performance of our method with that of the random walker [7], a pretrained convolutional neural network [9], the random forest pixel classifier [10], the power watershed [11], and band pass segmentation [12].

The random walker (RW) can be thought of as a special case of our method when *κ* = 0, and is automatically generated as the first step of our segmentation algorithm (see Section III-A and Fig. 1).

**Fig. 1.**
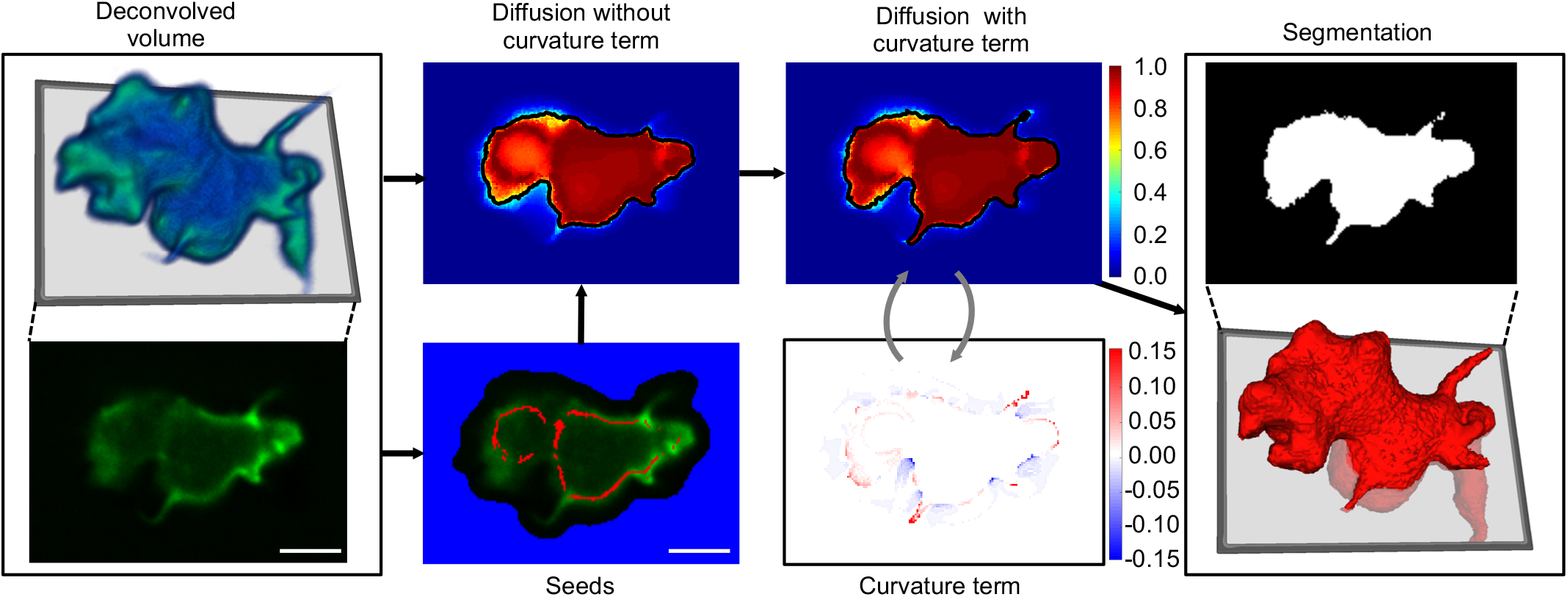
Flow chart outlining the major steps in the segmentation algorithm. *Dictyostelium* cells expressing LifeAct-GFP were imaged on an iSPIM microscope, as described in Section III-D3. 3D image pseudocoloured by intensity (blue represents low intensity, green represents high). Automatic seed selection for foreground (red) and background (blue) is based on Phansalkar thresholding [15], as described in Section III-B. The random walker method is implemented by simulating diffusion until equilibrium was reached, using the automatically selected seeds and image gradient-based weighting, as described in section III-A. Diffusion with an additional curvature term is subsequently simulated, with the same inputs as before, using the equilibrium values of the previous simulation as initial conditions, as described in Section III-A. Contour lines show the location of the 0.5 isosurface, which corresponds to the segmentation boundary. The curvature term is dependent on the diffusion system, and is periodically updated and fed back into the diffusion system. The curvature term is constructed to be strongest close to the segmentation boundary, and restricted to negative values outside this boundary and positive values inside. Scale bars represent 5 *μ*m.

**Fig. 2.**
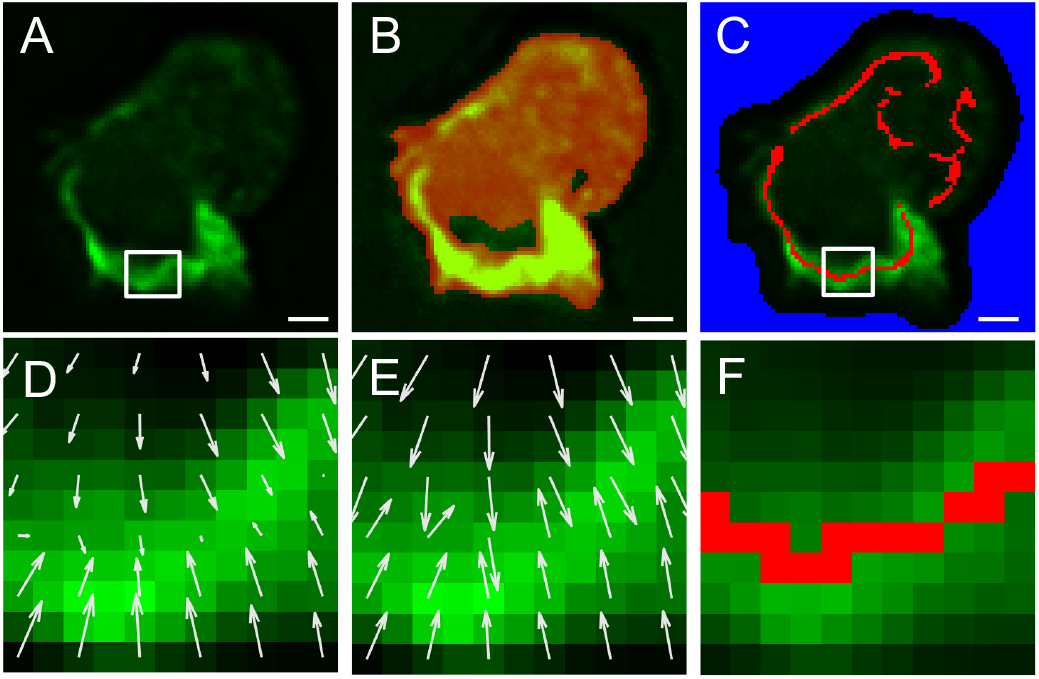
Automated seed selection. A: A slice through a volume to be segmented. White box indicates the position of D–F. B: The result of applying the Phansalkar threshold (red channel). C: Automatically selected background (blue) and foreground (red) seeds. White box indicates the position of F. Foreground seeds are obtained from the thresholded points according to D–F. D: Gradients, subsampled for ease of visualization. Gradient magnitude is used to identify local maxima. E: Normalized maximal gradient directions, used to compare gradient magnitudes. F: Foreground seeds resulting from comparing gradient magnitudes (D) along the locally maximal gradient direction (E). Scale bars represent 2 *μ*m.

We compared our method to the pretrained neural network published by Castilla *et al.* [9] (CNN). This network was trained on real and synthetic images of cells with multiple filopodia and therefore should be a close match for our data. In particular, this network was trained, in part, using synthetic datasets produced in the same manner as those from the Cell Tracking Challenge [12], [13] described below.

The random forest pixel classifier (RF) was implemented using the fast random forest method in the Trainable Weka Segmentation 3D plugin [53] for Fiji [54]. Because this classifier determines foreground and background voxels based on local voxel data, the automated seeding developed for our segmentation method was inappropriate to use for training due to it selecting only local maxima as foreground. For this reason, we used random sampling of the ground truth foreground and background to train the classifier. We chose sampling rates *N* of 50 and 500 training samples per slide for our comparison; a rate of *N* = 50 samples per slide is comparable to the number of automatically selected foreground seeds in Section III-B, while the higher rate of *N* = 500 was chosen to represent a highly fitted classifier. Once trained, the classifier was applied to the whole image stack to select foreground and background voxels. Based on preliminary testing, voxels were classified using the Gaussian, Hessian, and Laplacian features [53].

The power watershed (PW) is given by the energy optimization of Eq. 1 with finite *q* and *p* → *∞* [11]. In the following, we use a value of *q* = 2 as used by Couprie *et al.* [11]. This method is also related to the random walker, in that it represents the limiting case of *β* → *∞*.

The band pass segmentation method (BP) uses two pre-processing steps. The image stack *I* is initially clipped by a maximal intensity *I*_max_ to yield 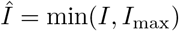. This image is subsequently smoothed using 3D Gaussian filters of standard deviation *σ_B_* and *σ*_*S*_, to give two smoothed images *I_B_* and *I_S_*. For image volumes with anisotropic resolution, *σ_B_* and *σ_S_* are scaled down in *z* by the ratio of the slice spacing to planar resolution. These images are combined to give the band pass image *I_BP_* = *I_S_ − αI_B_* for some scaling parameter *α*. Finally, a threshold *τ* is applied to *I_BP_*, followed by a morphological fill operator to obtain the final segmentation. The parameters *I*_max_, *σ_B_*, *σ_S_*, *α*, and *τ* were optimized using the coordinate ascent method for each dataset as described in the original formulation [12]. Parameter values for the band pass segmentation of the synthetic shape and real microscopy data are given in the supplementary materials.

### D. Data acquisition and pre-processing

#### 1) Synthetic test image

We constructed a synthetic test image to evaluate segmentation of protrusions and invaginations. Full details of how the image was generated can be found in the supplementary material. Briefly, a shape was generated with multiple protrusions and invaginations of varying widths (Fig. 4, Supplementary Fig. 2A). The corresponding binary image volume was blurred and Poisson noise was added to create a boundary texture similar to that of real cells (Supplementary Fig. 2C). The plane *z* = 0 was taken as the foreground seed set (the bottom of the shape in Supplementary Fig. 2A), while background seeds were taken to be the plane *z* = 80 (above the shape in Supplementary Fig. 2A).

#### 2) Cell Tracking Challenge images

We tested our method on Cell Tracking Challenge datasets Fluo-C3DH-A549 and Fluo-C3DH-A549-SIM (www.celltrackingchallenge.net) [12], [13]. These datasets contain real and simulated 3D confocal images of lung cancer cells, respectively. The simulated images were generated with the FiloGen model-based cell generator [20], [21]. Both datasets have planar pixel width of 0.27 *μ*m and slice separation 2.3 *μ*m. Test and training datasets are publicly available from the challenge website, but ground truth segmentations are only available for the training datasets. Our results for the test datasets were externally validated by the Cell Tracking Challenge, which allows a comparison of our results with other state of the art methods in Section IV-B2. The measure used by the Cell Tracking Challenge is based on the Jaccard score, but includes a factor to measure cell detection [12]. The methods compared below all detected the cells perfectly, and therefore this measure is equivalent to the Jaccard score. Additionally, we use our own comparison measures on the training dataset of Fluo-C3DH-A549-SIM to compare our results with the methods presented in Section III-C.

Pre- and post-processing steps were required for the PW, RW, and CERW, and are detailed in the supplementary material. For all three methods, the pre-processing steps were rescaling in *z* to obtain isotropic resolution, applying a band pass filter, and applying 2D contrast-limited adaptive histogram equalization [55] to movie 02 in Fluo-C3DH-A549-SIM. In order to better capture the long and branching filopodia in movie 02 of both datasets, a high value of *κ* was used. Because the curvature term is independent of the original image, this led to the filopodia being detected with a wider cross section than desired. Additionally, a higher curvature weighting led to background leaking into the cell through areas of low membrane intensity in some images. Accordingly, the following postprocessing steps were used (see the supplementary material for more information). The first step was to apply morphological dilation, fill, and erosion to fill any holes inside the cell. The second step was to erode or dilate the segmented cell locally based on the gradient magnitude of the original image, using a surface mesh of the segmented cell. This was performed by considering the line along the direction normal to the surface at each vertex *v* within a distance *r* from the surface. The value of *r* was calculated locally to be the largest value below a fixed limit *r_max_* where the closest surface vertex to any point on the line is *v*. If the position of the maximum gradient magnitude along this line was inside the segmented shape, then all voxels along the line between this point and the surface were assigned background values. Otherwise these voxels were assigned foreground values.

Background seeds were generated as described in Section III-B. Foreground seeds for the first movies of both datasets were selected using the dilation-fill-erosion method described in Section III-B, while for the second movies of both datasets the union of the seed set from the dilation-fill-erosion method and the maximal surfaces method was taken as the foreground seed set. Parameter values for seed selection can be found in the supplementary material.

#### 3) Microscopy data

We tested our method on 11 microscopy image stacks from two separate time series that had been manually segmented by a single annotator for comparison. *Dictyostelium* cells expressing LifeAct-GFP were imaged on an inverted selective plane illumination microscope (iSPIM) [56]. The resulting image stacks had pixel width 0.165 *μ*m and slice thickness 0.2 *μ*m. These images were deconvolved using the Richardson-Lucy algorithm using 50 iterations [57]. Deconvolution allows for more precise segmentation by reducing image blur. Preliminary testing suggested that rescaling these image volumes to produce isotropic resolution yielded little change in the segmentation, and therefore the image volumes were treated as having isotropic resolution to decrease processing times. The lack of improvement from rescaling may be due to a higher level of blurring present in the *z*-direction after deconvolution. The image volumes selected for segmentation each contained a single cell undergoing macropinocytosis. For computing the RW and CERW, all microscopy image volumes were normalized to [0, 1], and gamma correction (0.5) was applied.

Manual segmentations were drawn onto the 2D planes of the image volumes, cycling through (*x, y*)- (*x, z*)-, and (*y, z*)- planes iteratively until a satisfactory segmentation was achieved. A full description of how this was performed in 3D Slicer (www.slicer.org) [58] is provided in the supplementary information. Due to the subjective nature of manual segmentation, 3 of these image volumes were manually segmented by an additional 2 annotators, and a majority voting method was used to generate a fourth set of annotations, as is common practice [9], [12]. Inter-annotator variability and comparisons of the segmentation results with each of these three alternative annotations are included in the supplementary information.

The segmentation results below all use manual segmentations from annotator 1 as ground truth. These scores are compared to the scores between annotator 1 and the other annotators (Supplementary Table V) to determine if a segmentation method is indistinguishable from a human annotator with respect to any of the evaluation methods given in Section III-E. Accordingly, for each evaluation measure, we define the minimum annotator agreement to be the worst score (lowest Jaccard or local Jaccard score, highest Hausdorff distance or boundary displacement error) between annotator 1 and the other annotators. All segmentations with a score better than this minimum annotator agreement (higher Jaccard or local Jaccard score, lower Hausdorff distance or boundary displacement error) are deemed to be of equal merit with respect to the given measure. In these cases, comparisons with the alternative annotations will be used to confirm that the segmentation scores show similar results when given alternative interpretations. All real cell images and manual segmentations can be found at https://pilip.lnx.warwick.ac.uk/TMI_2020/.

### E. Evaluation methods

The measures used to evaluate the segmentation results were the Jaccard score, a localized form of the Jaccard score, the Hausdorff distance, and the mean boundary displacement error [59]. The first two measures describe how a segmentation performs on a voxel-wise basis, while the second two measures describe how the surface of the segmented shape differs from that of the ground truth.

For a ground truth image stack with foreground voxel set *S_GT_* and segmented image stack with foreground voxel set *S_SEG_*, the Jaccard score for the segmentation is given as *Jac* = |*S*_*GT*_ ⋂ *S*_*SEG*_ / *S*_*GT*_ ⋃ *S*_*SEG*_|. Here, we find it more informative to compare segmentations to ground truth in a localized fashion. For a point *p* with neighborhood *N* (*p*), the local Jaccard score *Jac_loc_* is given as

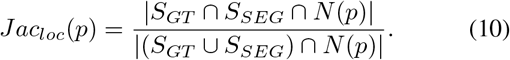

Because this measure is defined as a proportion of foreground voxels in a local neighborhood, points with a neighborhood populated with mostly foreground voxels will show less variation than a neighborhood populated with mostly background voxels. To address this difference, we also take the local Jaccard score of the negative spaces:

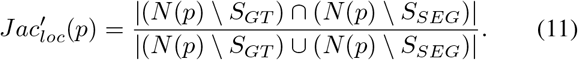

The overall local Jaccard score is thus defined as the lowest of these two scores:

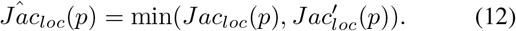

In the Cell Tracking Challenge and microscopy datasets, it is of particular interest to evaluate the segmentation at points on the cell surface of highly positive and negative curvature, because these correspond to biologically significant features. We therefore separate points on the surface of the ground truth into three sets: points with highly negative mean curvature (*H* < −0.2), highly positive mean curvature (*H* > 0.2), and points with low absolute curvature (|*H*| ≤ 0.2). Areas of highly negative mean curvature (denoted by *H*^−^) correspond to macropinocytotic cups, while areas of highly positive mean curvature (denoted by *H*^+^) correspond to filopodia.

For boundary difference measures, we first define the boundary of a shape in a binary image volume as the set of foreground voxels with at least one adjacent background voxel (26-connected). The distance from a point *p* to a boundary set *B* is given as

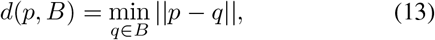

where ||·|| is the Euclidean norm. For a ground truth boundary set *B_GT_* and segmented boundary set *B_SEG_*, the Hausdorff distance is given as

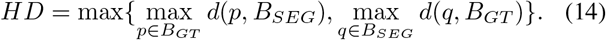

The mean boundary displacement error (BDE) is an asymmetric measure of the distance between the surfaces. The mean BDE from *B_GT_* to *B_SEG_* is given as

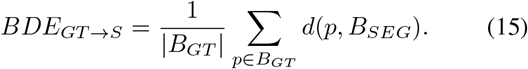

The mean BDE from *B_SEG_* to *B_GT_* is similarly defined. The Hausdorff measure reflects the area on the surface where the segmentation performs the worst, and a high Hausdorff distance can indicate failure to detect either protrusions or invaginations. The mean BDE gives a more directed measure, where a high BDE from ground truth to segmentation would indicate a failure to capture protrusions, while a high BDE in the reverse direction would indicate a failure to capture invaginations.

## IV. Results

We tested the curvature-enhanced random walker (CERW), random walker (RW), pretrained convolutional neural network (CNN), random forest pixel classifier (RF), band pass segmentation (BP), and power watershed (PW) on a synthetic 3D test image, data taken from the Cell Tracking Challenge dataset (www.celltrackingchallenge.net) [12], [13], and our own microscopy data. Four evaluation methods were used to compare the performance of these segmentation methods: the Jaccard score, the localized Jaccard score, the Hausdorff distance, and boundary displacement error. Our method was also externally evaluated by the Cell Tracking Challenge.

### A. Synthetic test image

#### 1) Relationship between β and κ

The first experiment using the synthetic test shape aims to investigate the relationship between *β* and *κ* in terms of segmentation performance. Here we use the localized Jaccard score with a neighborhood of square cross-section (width 21) and depth equal to the depth of the stack (80 slices). This neighborhood was constructed to include the full lengths of any protrusions or invaginations that intersect with the neighborhood. Here we investigate the minimum local Jaccard score *J*_min_, which highlights the score corresponding to the area of the shape where the segmentation performed the weakest. The results of this experiment are shown in Fig. 3. This shows that the curvature enhancement improves the random walker segmentation for all values of *β*. Additionally, we see that the values of *β* and *κ* that optimize the local Jaccard score are negatively correlated.

**Fig. 3.**
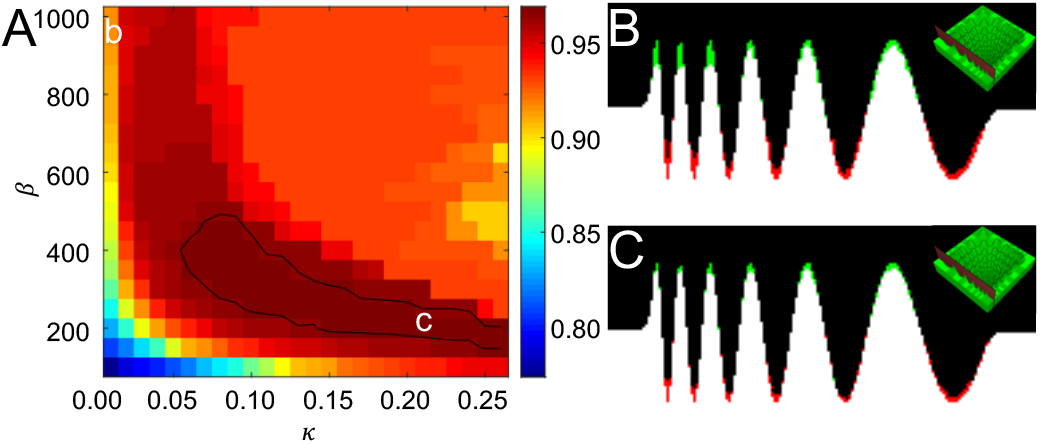
Results of experiments on the synthetic test image. A: Minimum local Jaccard scores (Eq. 10) for a range of parameter values. The optimal values of *β* and *κ* are negatively correlated, although even at high values of *β* a non-zero value of *κ* shows improvement on the standard random walker. Black contour line represents the 90th percentile. B: Comparison of the segmentation algorithm at point b (standard random walker) with the original mask. C: Comparison of the segmentation algorithm at point c (optimal values of *β* and *κ*) with the original mask. Red represents false positives, green represents false negatives, inset shows the position of the slice.

**Fig. 4.**
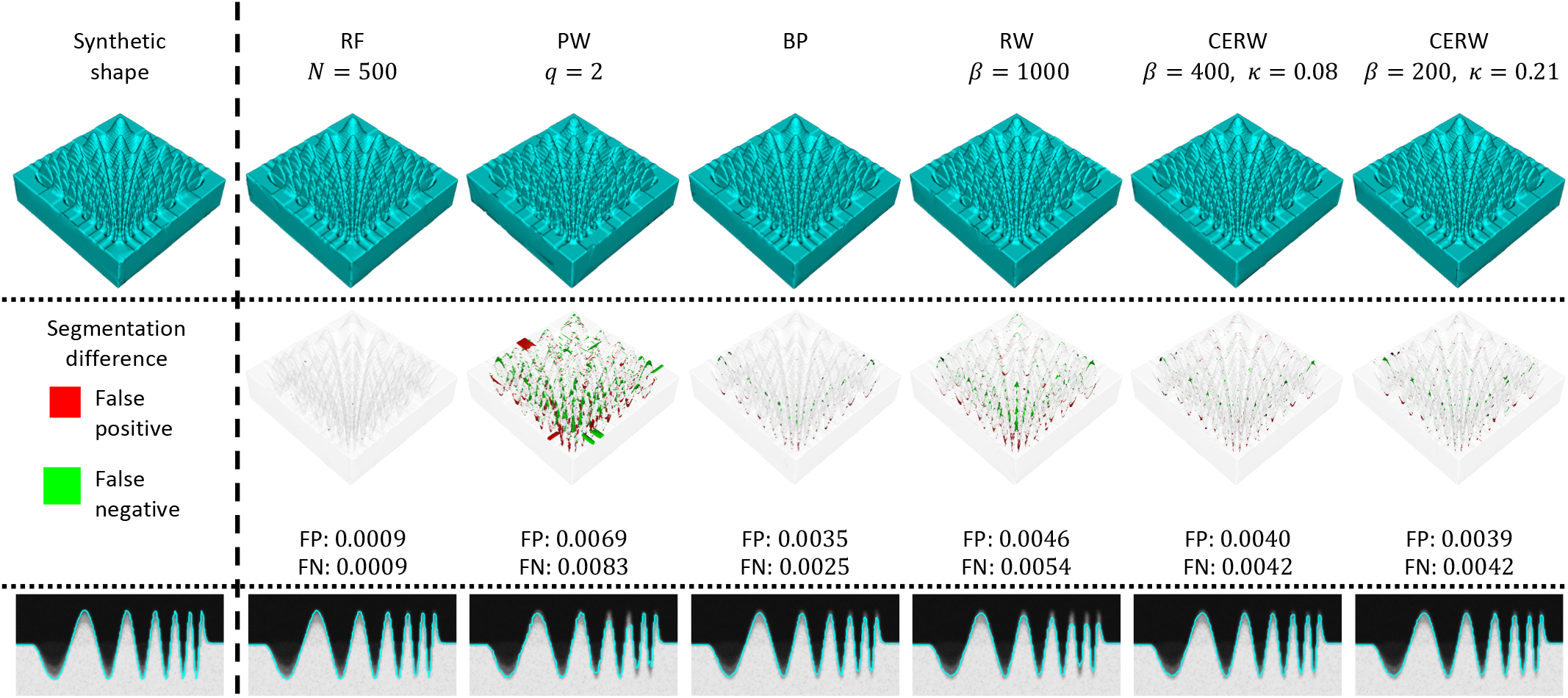
Comparison between the synthetic shape mask and random forest classifier (RF), power watershed (PW), band pass segmentation (BP), random walker (RW), and curvature-enhanced random walker (CERW). Top row: synthetic shape surface (left) and surfaces generated by each segmentation method. Middle row: differences between segmentation results and the synthetic mask, where red marks false positive and green marks false negative. FP: false positive rate, FN: false negative rate. Bottom row: slices through each surface compared to the original image volume. While the RF clearly performs best, this is likely due to the synthetic nature of the image. The BP shows better results than the remaining methods, and the CERW outperforms both PW and RW, only showing small errors for both parameter sets.

#### 2) Comparison with other methods

The broader set of segmentation methods are compared in Table I and Fig. 4. Unsurprisingly, the RF outperforms all other methods in this example, due to the fact that the image being analyzed is directly based on a binary image. Of the remaining methods, either the CERW or BP perform best, depending on the measure. More specifically, the BP performs best in the global measures (Jaccard score and mean BDE), while the CERW performs best in local Jaccard score, and the methods are tied on the Hausdorff distance. This suggests that the BP is able to recover more of the overall shape than the CERW, but loses some of the finer details of the shape, which are better-identified by the CERW. The CERW performs better than both the RW and PW in all scores. The PW and, to a lesser extent, the RW are particularly affected by the noise added to the image, as can be seen in Fig. 4. This noise sensitivity is avoided in the CERW because it enables the value of *β* to be reduced (compared to the optimal RW), which reduces the impact of variable gradients between neighboring voxels. Additionally, the curvature enhancement is able to compensate for the increased impact of blurring in the finer features, as can be seen in the slices in Fig. 4. Note that the CNN failed to segment the synthetic shape due to the marked difference to the training data and has been omitted from these results.

**TABLE I.** Comparison of Segmentation Methods Applied to The Synthetic Image.

| Method<sup>a</sup> | Jaccard | $J_{\min}$ | $H^b$ | Mean BDE<sup>c</sup> | |
| --- | --- | --- | --- | --- | --- |
|  |  |  |  | S→GT | GT→S |
| RF ($N = 50$) | 0.997 | 0.979 | 6.00 | 0.019 | 0.044 |
| RF ($N = 500$) | **0.998** | **0.987** | **5.10** | **0.012** | **0.022** |
| PW ($q = 2$) | 0.985 | 0.922 | 8.54 | 0.104 | 0.194 |
| BP | *0.994* | 0.955 | *8.00* | *0.041* | *0.092* |
| RW ($\beta = 1000$) | 0.990 | 0.917 | 13.0 | 0.065 | 0.162 |
| CERW (1000, 0.05) | 0.991 | 0.959 | 13.0 | 0.068 | 0.113 |
| CERW (400, 0.08) | 0.992 | 0.969 | *8.00* | 0.061 | 0.108 |
| CERW (200, 0.21) | 0.992 | *0.969* | *8.00* | 0.061 | 0.108 |

<sup>a</sup> RF: random forest classifier; PW: power watershed; BP: band pass segmentation; RW: random walker; CERW: curvature-enhanced random walker.
<sup>b</sup> Hausdorff distance
<sup>c</sup> Boundary displacement error. S→GT: mean distance from segmentation to ground truth; GT→S: mean distance from ground truth to segmentation.
Bold numbers highlight the best value, italic numbers highlight the best non-classifier value.

### B. Cell Tracking Challenge data

#### 1) Optimization of β and κ

Using the information gained in Section IV-A1 on the relationship between *β* and *κ*, we employed an optimization strategy of initially obtaining the optimal value of *β*, and subsequently optimizing kappa for a range of *β* values below this optimal *β*. Due to the large processing times (see Section IV-D), the optimization method was coarse-grained, and may be improved on with further processing. The optimal value of *β* for *κ* = 0 was identified by incrementing the value by 500 from an initial manually selected estimate until the Jaccard score for the RW reached a maximal value. The equilibrium values from these incremental steps were subsequently used as the initial conditions for the CERW. Starting with *β* at the optimal value for *κ* = 0, the optimal value of *κ* was identified by incrementing *κ* by 0.1 until a value producing the minimal Hausdorff distance was found, subject to a minimum Jaccard score of 0.6. The value of *β* was then decreased by 500 and the same method was used to optimize *κ*, starting at the optimal value of *κ* from the previous step. This sequence of decreasing *β* by 500 and optimizing *κ* was repeated for all values of *β* used in the initial optimization of *β*.

#### 2) Comparison with other methods

The results of the CERW and the two other highest-performing segmentation methods competing in the Cell Tracking Challenge are shown in Table II. Our method attained second place in Fluo-C3DH-A549 (real cells) and third place in Fluo-C3DH-A549-SIM (simulated cells). Both of the other high-scoring methods utilize CNNs and are therefore dependent on training data. In particular our method performs almost as well as the highest performing CNN in real cells. This result is in part due the fact that the CERW appears to perform better in the real cell images, but also may be due to the reduced amount of training data available for this dataset. Since our method requires no training data, it has an advantage in segmenting cells where manually annotated ground truth is difficult to obtain.

**TABLE II.**
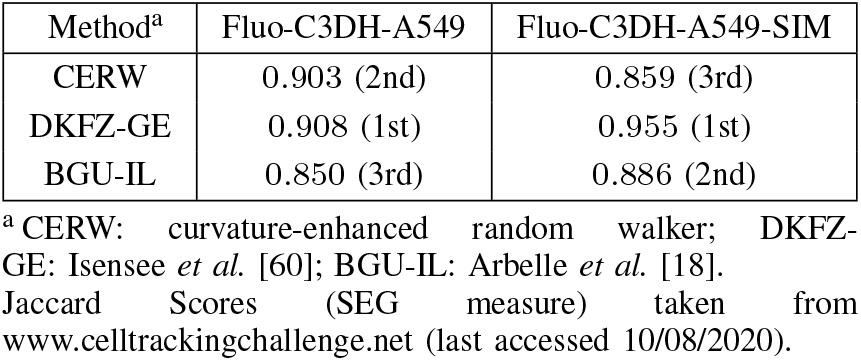
Cell Tracking Challenge results

| Method<sup>a</sup> | Fluo-C3DH-A549 | Fluo-C3DH-A549-SIM |
| --- | --- | --- |
| CERW | 0.903 (2nd) | 0.859 (3rd) |
| DKFZ-GE | 0.908 (1st) | 0.955 (1st) |
| BGU-IL | 0.850 (3rd) | 0.886 (2nd) |

<sup>a</sup>CERW: curvature-enhanced random walker; DKFZ-GE: Isensee *et al.* [60]; BGU-IL: Arbellet *et al.* [18]. Jaccard Scores (SEG measure) taken from www.celltrackingchallenge.net (last accessed 10/08/2020).

We use our own comparison measures on the segmentation of training data from Fluo-C3DH-A549-SIM in order to gain a more detailed understanding of the performance of the CERW. For the purposes of comparison, the binary image stacks for ground truth, RF segmentation, and BP segmentation were scaled in *z* to achieve isotropic resolution, using the nearest neighbors method for interpolation. We use the localized Jaccard score with neighborhood given by a cube of side 21 voxels (5.67 *μ*m), evaluated at all points on the simulated cell surface. Curvature is used to separate this measure into areas of highly negative (*H*^−^), low (*H*^0^), and highly positive (*H*^+^) mean curvature, as described in Section III-E. Of particular interest in this dataset is the accuracy of filopodia segmentation, which is most strongly reflected in the local Jaccard score in *H*^+^ areas, the mean boundary displacement error from ground truth to segmentation (*BDE*_*GT→S*_), and the Hausdorff distance. We focus our attention on the CNN, RW, and CERW, which are the best-performing machine learning and pixel-based methods, respectively, in both movies. The results for these methods can be found in Table III, while the full results are summarized in Supplementary Table IV.

**TABLE III.**
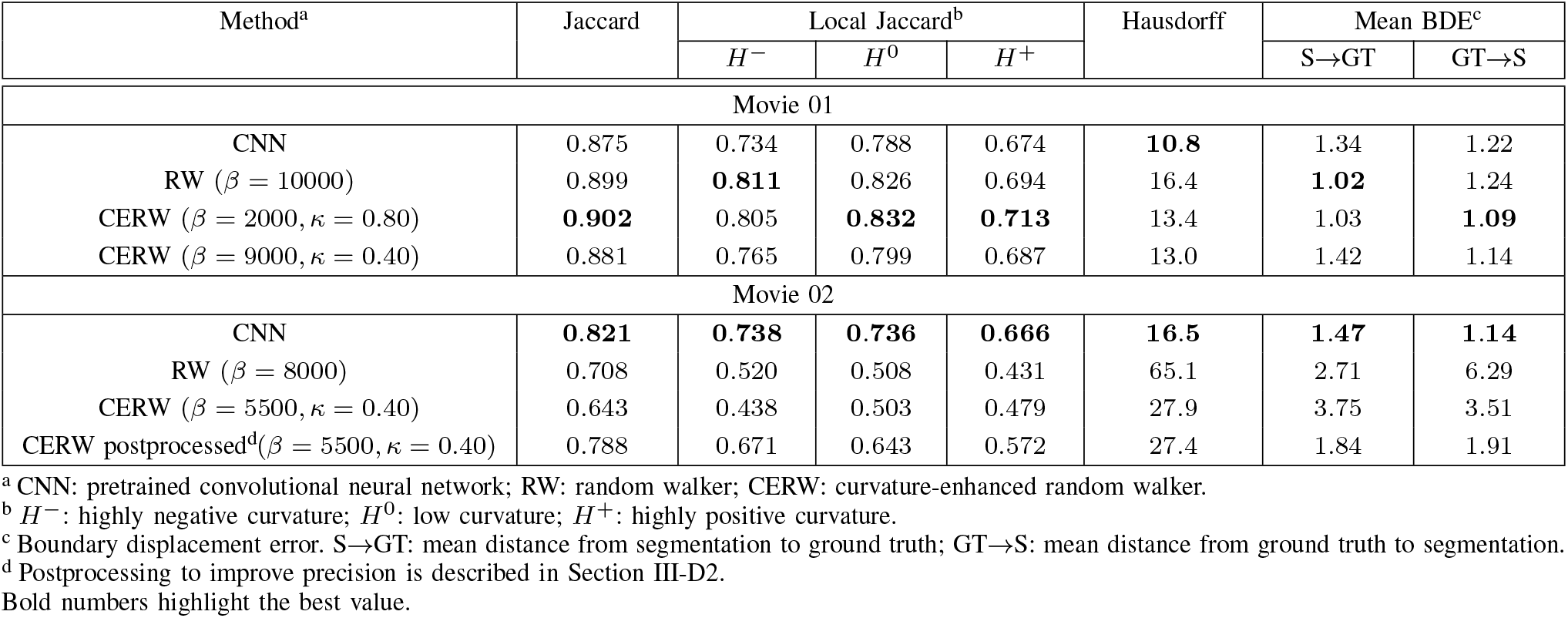
Comparison of Segmentation Methods Applied to Cell Tracking Challenge Data

In the first movie, the RW and CERW outperform all other methods in all scores except the Hausdorff measure, where the CNN performs best. The curvature enhancement yields improvements in measures corresponding to protrusions (loc Jaccard in *H*^+^ areas, Hausdorff, and *BDE*_*GT→S*_) and gener evaluation measures (Jaccard and local Jaccard in *H*^0^ area while decreasing accuracy in other measures. This sugge that, while the curvature enhancement shows an improveme in filopodia detection, as can be seen in the top row of Fig. there are other areas of the cell where the curvature enhanc ment decreases the accuracy of the segmentation. the CERW generally outperforms the CNN, which was train on similar datasets [9]. The top row of Fig. 5 shows that, whi the CNN detects the position filopodia well, it produces long filopodia than in the ground truth, an effect not present in t CERW. This suggests that the CERW can improve upon de learning-based methods even with an abundance of traini data.

**Fig. 5.**
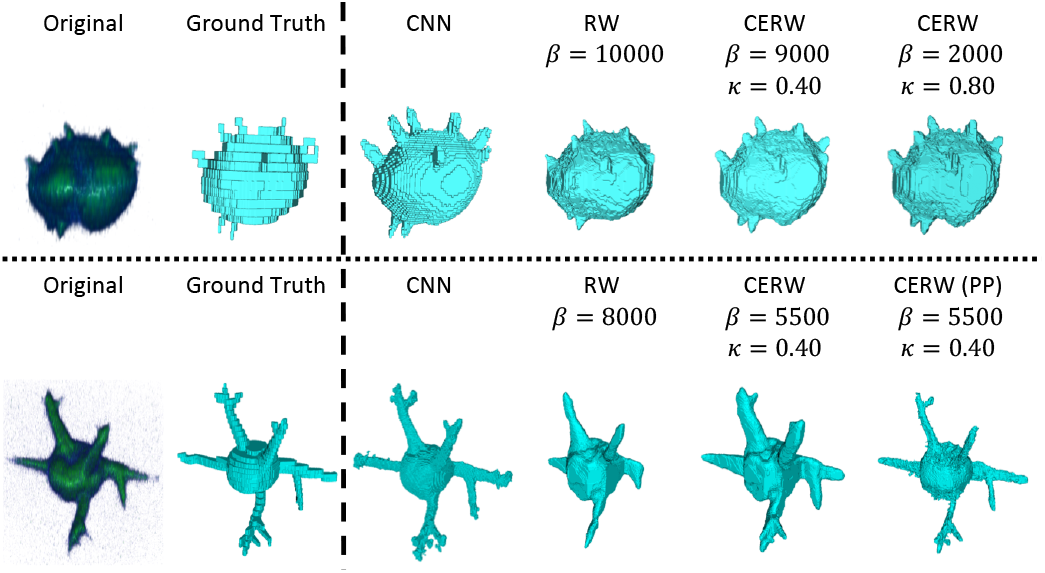
Comparison between Cell Tracking Challenge ground truth and pretrained convolutional neural network (CNN), random walker (RW), and curvature-enhanced random walker (CERW). Top row: surfaces of each segmentation for movie 01. Bottom row: surfaces of each segmentation for movie 02. In the first movie, the CERW better captures the full length of the filopodia than the RW, while the CNN segmentation gives filopodia longer than in the ground truth. The second movie shows that the CERW is able to recover filopodia branches not identified by the RW, identifying most of the branching structures. The post-processing step described in Section III-D2 reduces the width of the filopodia, but has little impact on the length. The CNN is better able to capture the full extent of these branching structures, although again some length has been added in comparison to the ground truth. Binary volumes for ground truth have been rescaled using the nearest neighbors method, which leads to the step effect in the images.

The second movie demonstrates how the curvature enhanc ment can dramatically improve the random walker segment tion. In Fig. 5, it is clear that there are several branching stru tures that are absent in the RW segmentation, but have been recovered by the CERW. In comparison with other metho the CNN performs best in all measures, as expected given that it was trained on similar datasets [9]. As shown in Fig. 5, the CNN is better able to detect the branches of the filopodia th the CERW. A limitation of the CERW is that while larger values of *κ* could potentially improve the segmentation of these branching structures, this will destabilize the system. This issue could be resolved by spatially varying *κ*, but this extension is beyond the scope of this paper.

### C. Microscopy data

#### 1) Relationship between β and κ

As with the synthetic shape, we first evaluate how the segmentation is affected by the choice of *β* and *κ*. Here we use the localized Jaccard score with neighborhood given by a cube of side 21 voxels (3.465 *μ*m in-plane, 4.2 *μ*m in *z*) evaluated at all points on the manually segmented cell surface. We use local curvature to separate this measure into areas of highly negative (*H*^−^), low (*H*^0^), and highly positive (*H*^+^) mean curvature, as described in Section III-E. Of particular interest in these datasets is the accuracy of filopodia and cup segmentation, which are most strongly reflected in the local Jaccard scores in *H*^+^ and *H*^−^ areas, respectively.

The mean local Jaccard scores for a range of *β* and *κ* values for each curvature range are summarized in the top row of Fig. 6. As with the synthetic shape, these images show that the addition of the curvature enhancement improves the segmentation scores for all values of *β*. All three images show a negative correlation for the optimal values *β* and *κ*, but with the optimal range of values being offset depending on the curvature of the surface. The global Jaccard scores for each of the volumes are summarized in the bottom row of Fig. 6. Here we see a similar relationship between the optimized values of *β* and *κ* as observed in the localized scores and synthetic image. Examples of segmentations of all real image volumes are provided in the supplementary material.

**Fig. 6.**
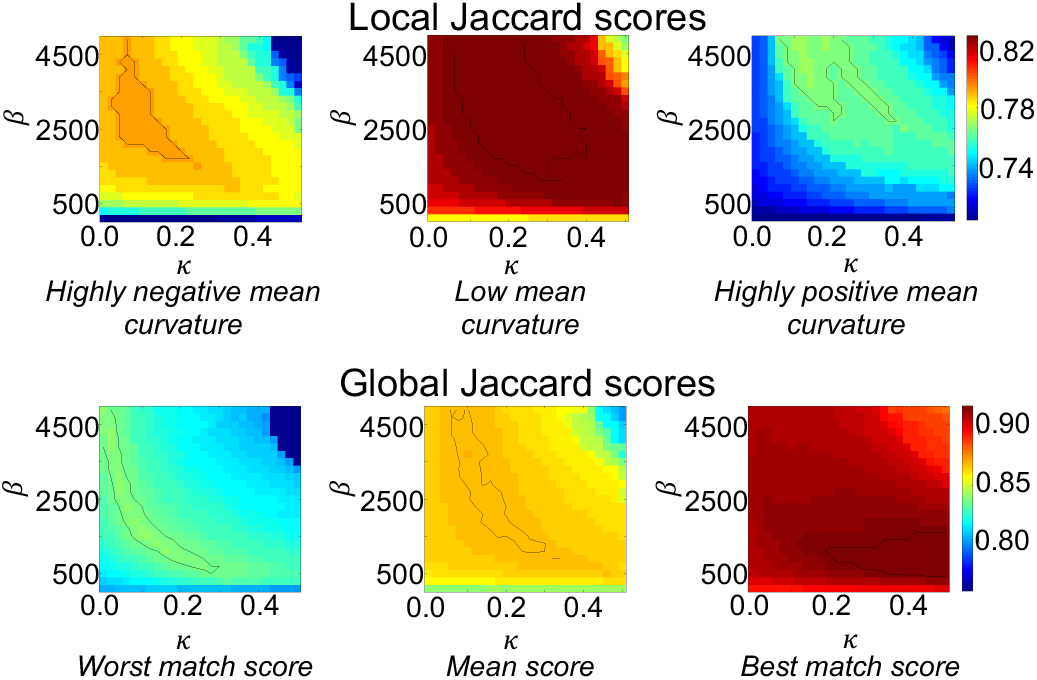
*Top row*: Mean local Jaccard scores for negative, low and positive curvatures of the surface of the manual segmentation of a cell, for a range of parameter values. As with the synthetic image, the optimal values of *β* and *κ* are negatively correlated. *Bottom row*: Minimum, mean, and maximum Jaccard scores for all segmented microscopy images for a range of parameter values. Again the negative correlation of the optimal values of *β* and *κ* is evident. Black contour lines represent the 90th percentile.

Comparing results in Fig. 6 to the minimum annotator agreement (Supplementary Table V), we see that the local Jaccard scores for areas of highly negative (Fig. 6 top left) and low curvature (Fig. 6 top middle) almost entirely lie above the minimum annotator agreement (0.744 and 0.784 respectively). This suggests that the addition of the curvature enhancement may not be producing a result that is any closer to the true segmentation in areas of highly negative or low curvature. However, for areas of highly positive curvature (Fig. 6 top right), all RW segmentations yield scores below the minimum annotator agreement (0.740). This suggests that the addition of the curvature term does produce results closer to the true segmentation in areas of highly positive curvature. Furthermore, the curvature enhancement does show improvement on the standard random walker for all measures in comparison to all other manual annotations (Supplementary Tables IX–XI).

#### 2) Comparison with other methods

Comparison of the CERW with other segmentation methods is summarized in Table IV. Of particular interest here are the scores corresponding to protrusions (local Jaccard in *H*^+^ areas, *BDE*_*GT→S*_, Hausdorff) and invaginations (local Jaccard in *H*^−^ areas, *BDE*_*S→GT*_, Hausdorff) since these correspond to the biological features of interest.

**TABLE IV.**
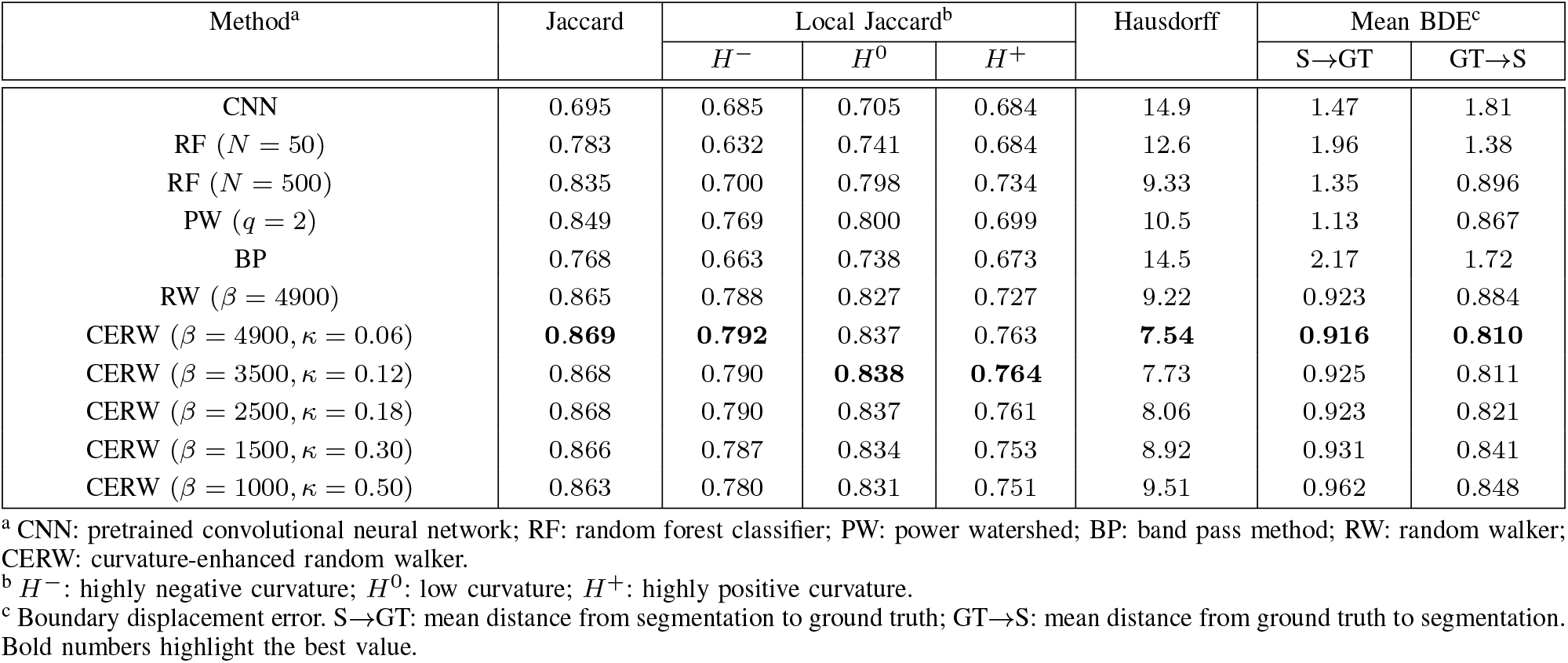
Comparison of Segmentation Methods Applied to Real Data.

| Method<sup>a</sup> | Jaccard | Local Jaccard<sup>b</sup> |  |  | Hausdorff | Mean BDE<sup>c</sup> |  |
| --- | --- | --- | --- | --- | --- | --- | --- |
| | | $H^-$ | $H^0$ | $H^+$ | | S→GT | GT→S |
| CNN | 0.695 | 0.685 | 0.705 | 0.684 | 14.9 | 1.47 | 1.81 |
| RF ($N = 50$) | 0.783 | 0.632 | 0.741 | 0.684 | 12.6 | 1.96 | 1.38 |
| RF ($N = 500$) | 0.835 | 0.700 | 0.798 | 0.734 | 9.33 | 1.35 | 0.896 |
| PW ($q = 2$) | 0.849 | 0.769 | 0.800 | 0.699 | 10.5 | 1.13 | 0.867 |
| BP | 0.768 | 0.663 | 0.738 | 0.673 | 14.5 | 2.17 | 1.72 |
| RW ($\beta = 4900$) | 0.865 | 0.788 | 0.827 | 0.727 | 9.22 | 0.923 | 0.884 |
| CERW ($\beta = 4900, \kappa = 0.06$) | **0.869** | **0.792** | 0.837 | 0.763 | **7.54** | **0.916** | **0.810** |
| CERW ($\beta = 3500, \kappa = 0.12$) | 0.868 | 0.790 | **0.838** | **0.764** | 7.73 | 0.925 | 0.811 |
| CERW ($\beta = 2500, \kappa = 0.18$) | 0.868 | 0.790 | 0.837 | 0.761 | 8.06 | 0.923 | 0.821 |
| CERW ($\beta = 1500, \kappa = 0.30$) | 0.866 | 0.787 | 0.834 | 0.753 | 8.92 | 0.931 | 0.841 |
| CERW ($\beta = 1000, \kappa = 0.50$) | 0.863 | 0.780 | 0.831 | 0.751 | 9.51 | 0.962 | 0.848 |

<sup>a</sup> CNN: pretrained convolutional neural network; RF: random forest classifier; PW: power watershed; BP: band pass method; RW: random walker; CERW: curvature-enhanced random walker.
<sup>b</sup> $H^-$ : highly negative curvature; $H^0$ : low curvature; $H^+$ : highly positive curvature.
<sup>c</sup> Boundary displacement error. S→GT: mean distance from segmentation to ground truth; GT→S: mean distance from ground truth to segmentation. Bold numbers highlight the best value.

The CERW outperforms all other methods in all scores. Fig. 7 shows this result in the context of an image volume (see also Supplementary Fig. 3). While the CNN performs well on seven of the images, it fails to identify much of the cell in the remaining four images. In the images where the CNN does perform well, the segmented filopodia tend to be longer than the ground truth, as can be seen in Fig. 7. The RF tends to have a low false negative rate but a high false positive rate, which manifests as an expanded boundary. Similarly, the BP has a high false positive rate, but also fails to identify all protrusions. The PW tends to be highly affected by noise, giving a rough boundary that can lead to large areas of false positives and large areas of false negatives. The RW provides a smoother segmentation, but fails to accurately segment the protrusions, due to variation in intensity along these protrusions (see slices in Fig. 7). The curvature enhancement allows the recovery of these protrusions, leading to a more accurate segmentation. Additionally, the false positive and false negative rates are much more balanced in the CERW than in the other methods, which suggests that it has less bias toward over- or under-segmenting the image.

**Fig. 7.**
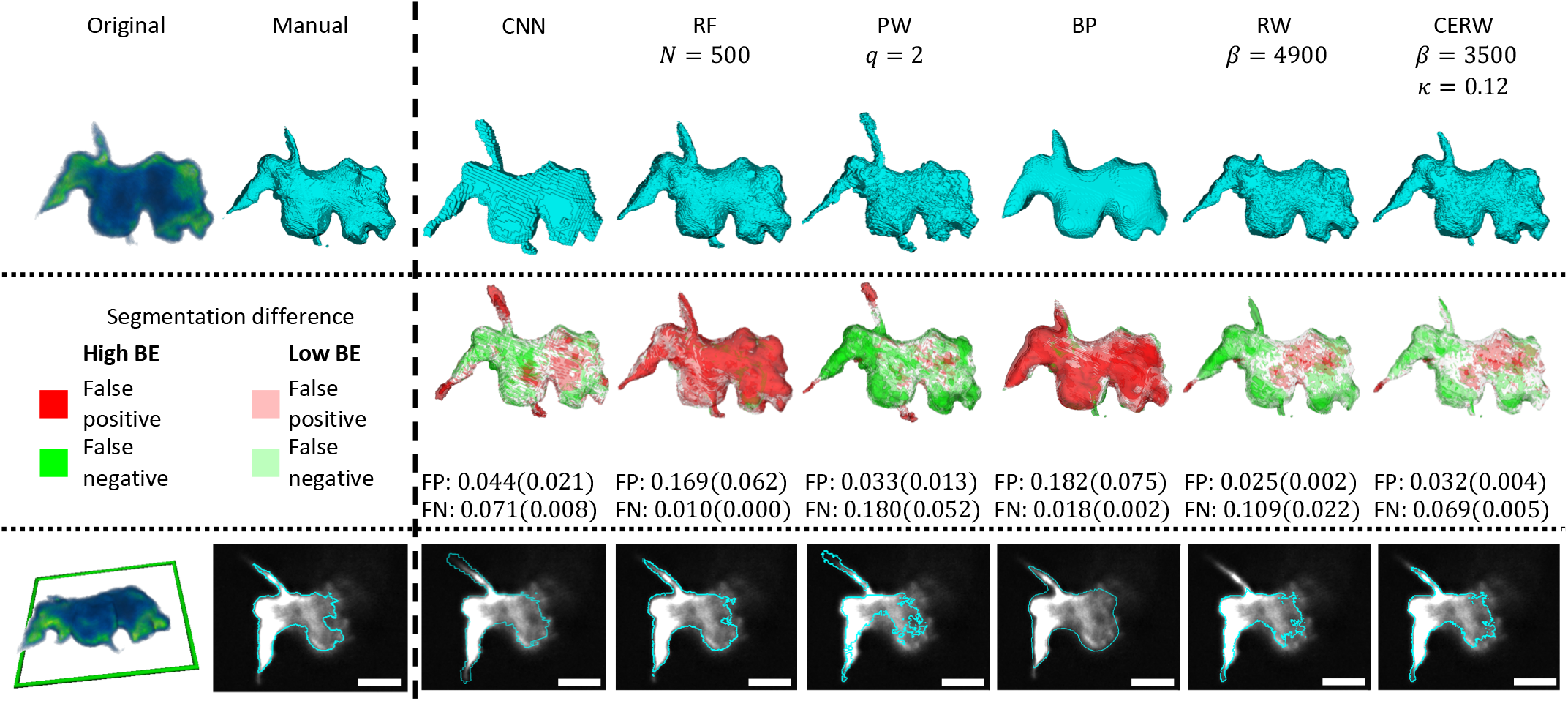
Comparison between manual segmentation and pretrained convolutional neural network (CNN), random forest classifier (RF), power watershed (PW), band pass segmentation (BP), random walker (RW), and curvature-enhanced random walker (CERW). Top row: surfaces of each segmentation. Middle row: differences between segmentation results and manual segmentation, where red marks false positive, green marks false negative, and reduced opacity is given to voxels with a boundary error of 1 (low BE). FP: false positive rate, FN: false negative rate, numbers in brackets are error rates of high BE points only. Bottom row: slices through each surface compared to original image, scale bars represent 5 *μ*m. The CERW outperforms all other methods, with almost all errors having low boundary error. Filopodia detected by all other methods are excessively long (CNN, RF, PW), short (BP, RW), or wide (RF, BP). Note a region on the left is marked as false positive in most segmentations. The slices in Supplementary Fig. 4 give alternative manual segmentations from separate annotators, showing that only one annotator has included this region in the cell, and that the majority vote excludes this region.

The CNN, RF, BP, and PW all yield scores worse than the minimum annotator agreement (Supplementary Table V). For most measures, both the RW and CERW score better than the minimum annotator agreement. This suggests that random walker-based methods yield segmentations closer to the true segmentation than all other methods. The measures where the RW performs worse than the minimum annotator agreement are the local Jaccard score in *H*^+^ areas (as observed in Section IV-C1) and the Hausdorff distance. These measures are both formulated to detect differences in protrusion detection, and therefore the improvement in these measures from the curvature enhancement indicates an improvement in detecting protrusions in the true segmentation.

Similar results to those shown in Table IV can be seen in comparison with the other manual segmentations and majority voting segmentations in Supplementary Tables IX–XI, with a notable exception that the CNN performs considerably better on the three images that were segmented by multiple annotators. Here we see that the CNN scores better than the CERW in comparison to the individual annotators in some scores, but worse in all scores in comparison with the aggregated segmentation, which suggests that the CERW provides a better segmentation than the CNN in these images. The other major difference in these tables in terms of comparative scores is that the RF performs better than our method in the Hausdorff distance. The RF scores worse than the CERW in all other measures, suggesting that the RF oversegments the image, yielding a good Hausdorff distance but poor precision, as can be seen in Fig. 7.

### D. Processing times

A comparison of the average time required to segment image volumes from each dataset with each method is provided in Supplementary Table XII. The CERW is the slowest method in all cases, with the larger and more complex image volumes in the second movie of the Cell Tracking Challenge data requiring roughly 20 times longer to process than the slowest non-random walker-based method (the PW). However, roughly half of the computation time for the CERW is taken up by the initial RW segmentation. We simulated diffusion for this step, in order to provide continuity between methods, but faster methods for computing the initial RW values are available [7], [8], which would greatly improve the speed. Furthermore, the method of simulation for the curvature enhancement step has not been optimized for time-efficiency, and could potentially be improved upon to reduce the segmentation time. Finally, the seed selection also requires a significant proportion of the processing time. However, this step was not performed on a GPU, which could significantly reduce computation time.

## V. Discussion

We have added a curvature term to the non-linear diffusion representation of the random walker method. This yielded an improvement to the original random walker in both real microscopy image stacks and the synthetic test image, and in filopodia detection in the Cell Tracking Challenge datasets. Furthermore, our method outperforms all other methods tested on our microscopy images, and performs competitively against state of the art deep learning-based methods in challenge datasets.

Our method is capable of segmenting cells with similar accuracy to deep learning-based methods even when those networks are trained with data captured in the same manner as the test data. Furthermore, our method is more reliable at segmenting our data than a pretrained CNN [9]. Since the CERW does not require manually annotated data with which to train, this result suggests that the CERW can be utilized to segment images where manual annotations are unavailable, such as those produced with novel markers or new imaging techniques. Additionally, not relying on manual annotations reduces the level of experimenter bias introduced into the results.

The main advantage of using the curvature-enhanced random walker over the standard random walker is the ability to recover finer details in the image, as is evident in Figs 5 and 7. This is especially useful in microscopy images, where image capture tends to blur these finer features and deconvolution can only partially compensate for this effect. In particular, given the recent discovery of the involvement of filopodia in macropinocytosis [3], being able to accurately segment these fine structures is vital for identifying the mechanisms involved.

Another advantage of the curvature-enhanced random walker over the standard random walker is that lower values of *β* can be used with a sufficiently large *κ*, which reduces the impact of noise on the segmentation. This is because a low value of *β* reduces the impact of noise-induced low-level fluctuations in image gradients, but also reduces the ability of the random walker to capture finer details. The introduction of a high level of curvature enhancement (large *κ*) enables the recovery of these details. Furthermore, since the curvature enhancement is independent of the original image, it is not affected by noise. This effect is especially advantageous when analyzing how the cell surface changes over time, since the segmentation surface is less prone to arbitrary noise-induced variations between time frames.

While we have not explicitly examined the stability of the curvature-enhanced random walker, our experiments show that there is a range of values of *β* and *κ* that produce stable results (see Figs 3 & 6). The only case where we observed instability was for high *β* and high *κ* (top right in Fig. 6). In such instances, the system did not converge, and this instability is only mitigated by the diffusion term in Eq. 7. This effect is likely due to the fact that the curvature enhancement term introduces instability into the system, which is only mitigated by the diffusion term in Eq. 7. If *β* is large, then this reduces the rate of diffusion, which in turn reduces the stability of the system. The question of stability will be a topic of future research.

The relationship between the optimal values of *β* and *κ* depends on the local curvature in our microscopy data (Fig. 6). Specifically, areas of highly positive curvature require a larger value of *κ* than areas of highly negative curvature to attain the optimal Jaccard scores, although there is a small overlap in these optimal values. Additionally, areas of highly positive curvature have a lower Jaccard score than areas of highly negative curvature for all seeded methods (see Table IV).

This is potentially due to the nature of the structures on the cell membrane that give rise to these curvature values; highly positive curvature corresponds to filopodia, which are long, thin protrusions on the cell membrane, while highly negative curvature corresponds to macropinocytotic cups, which in our data were much wider than the filopodia and relatively shallow. This means that highly positive curvature is associated with sharper features than highly negative curvature, which suggests that the variation in *κ* is linked to the shape of the features being segmented, rather than the sign of the curvature.

One major drawback of the current implementation of our method is the time required to process images, particularly the more complex images from the Cell Tracking Challenge. However, our implementation of the CERW only uses minimal speed optimization methods, and therefore we expect to dramatically improve on this issue in future implementations. One possible improvement would be to employ a faster random walker algorithm for the initial segmentation [8]. Improvements could be made by reducing the segmentation volume over time; for much of the image, the curvature enhancement has little impact, and therefore the curvature enhancement step could be restricted to areas where the curvature term is changing the segmentation. Finally, we note that our method is based on a PDE, and therefore could be computed faster by using more advanced PDE solvers.

Another limitation of our method is that in order to segment the large branching structures in Section IV-B2 a large value of *κ* is required, which can lead to segmentation errors in other areas of the cell. Furthermore, increasing *κ* beyond the level used leads to destabilization of the system. This issue could be addressed by varying *κ* based on the input image so that the curvature term only affects the relevant areas. This variation could also be used for the speed improvements mentioned above, by reducing the segmentation to areas with non-zero *κ*.

We have only investigated the application of our method to single-cell image stacks. The extension of the curvature-enhanced random walker to multi-cell image stacks can be applied in a manner similar to previous random walker implementations [7]. Our method could also be applied to 2D images, using curvature in place of mean curvature.

## VI. Conclusion and future work

We have shown that our curvature-enhanced random walker segmentation method yields improvements on the standard random walker method, and performs better than other state of the art methods in *Dictyostelium* image volumes. We have also shown that our method performs similarly to state of the art deep learning-based methods in externally evaluated data.

One limitation of our method is the time required to segment more complex image volumes. Future implementations will aim to deal with this drawback by using faster methods of simulation and seeding.

Extensions in the curvature enhancement could include spatially varying *κ*, or using a formulation to increase the Willmore energy of implicit surfaces. The curvature enhancement presented here could also be applied to other methods, such as active contour segmentation, regularized by shape priors to reduce instability [61].

In terms of biological data, we have only applied the curvature-enhanced random walker to light microscopy images of *Dictyostelium* and lung cancer cells. This method would also be well-suited to segmenting other objects with complex surface structures, such as computed tomography images of vertebrae or lungs, or electron microscopy images of complex cell structures.

## Supporting information

Manual Segmentation Protocol

## Acknowledgment

The authors would like to thank Rob Kay and Peggy Paschke (MRC-LMB Cambridge), for providing cell lines, the Warwick Advanced Bioimaging RTP for access to the DiSPIM microscope (BBSRC grant BB/M01228X/1), and the Cell Tracking Challenge organizers for permitting the use of the Fluo-C3DH-A549-SIM training data and externally evaluating our method.

