## Supplementary Material for "A Curvature-Enhanced Random Walker Segmentation Method for Detailed Capture of 3D Cell Surface Membranes"

### I. Supplementary methods

#### A. Seed selection

Automated seed selection is performed on microscopy data based on the Phansalkar threshold [1]. The Phansalkar threshold $T$ at a point $x$ is given by

$$T(x) = \mu(x)(1 + p\exp(-q\mu(x)) + k(\sigma(x)/r - 1)), \tag{1}$$

where $\mu(x)$ and $\sigma(x)$ are the local mean and standard deviation of the image intensity, and $p$, $q$, $k$, and $r$ are fixed parameters. The value of $p$ is set to 2, and the local neighborhood for calculating $\mu(x)$ and $\sigma(x)$ is given by a cube of side $w_{\text{phan}}$ voxels centered at $x$. The values of $q$, $k$, and $r$ are found based on each individual image volume. The value of $q$ is set so that the exponential decays by half over two standard deviations of the local means:

$$q = \log(2)/(2\sigma_\mu), \tag{2}$$

where $\sigma_\mu$ is the standard deviation of all local mean values in the image volume. The values of $r$ and $k$ are set based on the following assumptions. Foreground voxels are assumed to have local standard deviation equal to the maximum local standard deviation $\hat{\sigma}$, which maximizes $T$ in Eq. 1, and local mean

$$\mu_{FG} = \mu_\mu + 2\sigma_\mu, \tag{3}$$

where $\mu_\mu$ is the mean of all local means in the image volume, and $\sigma_\mu$ is as in Eq. 2. Conversely, background voxels are assumed to have zero local standard deviation (minimizing $T$ in Eq. 1) and local mean

$$\mu_{BG} = \mu_\mu + \sigma_\mu, \tag{4}$$

where $\mu_\mu$ and $\sigma_\mu$ are as above. We define $r$ and $k$ based on these assumptions:

$$r = \hat{\sigma}/(1 - \exp(-q(\mu_{FG} - \mu_{BG}))), \tag{5}$$

$$k = p(\exp(-q\mu_{BG}) + \exp(-q\mu_{FG})/(1 - \hat{\sigma}/r))/2. \tag{6}$$

These values ensure that any voxels conforming to the assumptions made on foreground voxels will have a threshold value below the local mean, while any voxels conforming to the assumptions made on background voxels will have a threshold value above the local mean. An example of the output of this thresholding is shown in Fig. 1B. Prior to thresholding, images were smoothed using mean and median filters to reduce the impact of noise on the seeding.

Background seeds are obtained by applying a spherical dilation operator of radius $r^d_{BG}$ voxels to the binary volume obtained by the Phansalkar thresholding, filling holes, applying a spherical erosion operator of radius $r^e_{BG}$, and inverting the resulting binary image volume. An example of the result of this process is shown in Fig. 1C.

Two methods are used for selecting foreground seeds for the data presented in Section III-D. The first is the same method as for the background seed selection, without the final inversion, with dilation and erosion radii being $r^d_{FG}$ and $r^e_{FG}$ respectively. In the second method, foreground seeds are selected to represent maximal surfaces in the image volume. Gradients are computed using a Sobel operator (Fig. 1D). Each voxel is assigned the direction of the largest magnitude gradient within a 26-connected neighborhood for the purposes of comparison (Fig 1E). Magnitudes along the assigned direction are linearly interpolated from the neighboring voxels, and a voxel is labeled as a turning point if its gradient magnitude is lower than these interpolated magnitudes. Turning points with a negative Laplacian are labeled maximal. This set of maximal points is restricted to points marked as foreground by the Phansalkar thresholding algorithm. To further improve the robustness of this algorithm, the maximal points are grouped into (6-)connected components. The largest connected component is labeled as foreground. Any other connected components larger than one voxel with mean intensity greater than the mean intensity of the largest connected component are also labeled as foreground. An example of the result of this process is shown in Fig. 1C. Parameter values for seed selection in all datasets are available in the Supplementary Table I.

As a result of this seeding method, the edges of the image volume tend to be populated with background seeds, meaning that the volume to be segmented is generally smaller than the input image volume. Accordingly, we automatically crop all images to the smallest bounding cube with background seeds populating the edge planes, in order to reduce processing times.

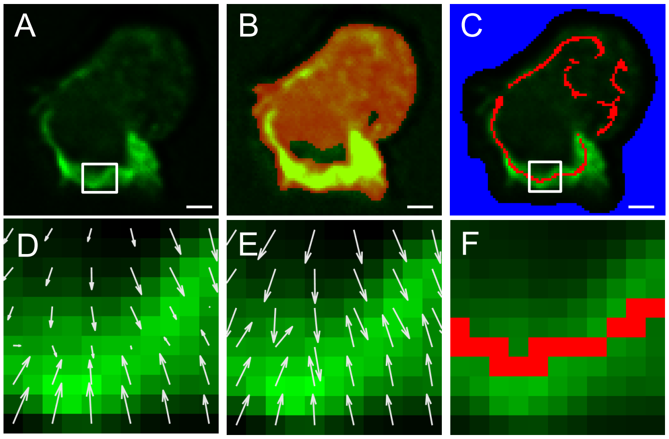


Supplementary Fig. 1. Automated seed selection. A: A slice through a volume to be segmented. White box indicates the position of D–F. B: The result of applying the Phansalkar threshold (red channel). C: Automatically selected background (blue) and foreground (red) seeds. White box indicates the position of F. Foreground seeds are obtained from the thresholded points according to D–F. D: Gradients, subsampled for ease of visualization. Gradient magnitude is used to identify local maxima. E: Normalized maximal gradient directions, used to compare gradient magnitudes. F: Foreground seeds resulting from comparing gradient magnitudes (D) along the locally maximal gradient direction (E). Scale bars represent 2 $\mu$m.

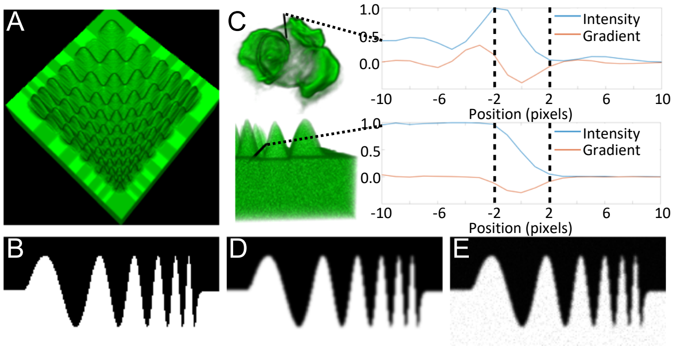


Supplementary Fig. 2. Generating the 3D synthetic test image. A: Synthetic shape created to test the effectiveness of the curvature-enhanced random walker algorithm. B: Slice through the synthetic shape. C: Normalized line scans of real and synthetic images. Comparison of these line scans informed the choice of parameter values for Gaussian blurring (D) and applying Poisson noise (E).

### *B. Synthetic test image construction*

Here we have chosen a shape with multiple protrusions and invaginations of varying widths (Fig. 2A). The resolution of the synthetic shape (Fig. 2B) was selected based on observation of microscopy data, such that the finest structures are only $1-2$ voxels wide. The intensity values were generated such that the shape boundaries have a similar pattern of intensities to that of real cells (Fig. 2C). The intensity pattern at a point on the manually identified boundary of a real cell was taken to be the voxel intensities along the line normal to the boundary. The pattern of intensities at the synthetic shape boundary was determined in the same manner. These line scans were normalized to $[0, 1]$, so that the gradients at the boundary could be compared up to a scaling factor. The aim in the following is to match the distribution of the steepest gradients of these line scans in the synthetic test image volume $\{g_s\}$ with that of the real data $\{g_r\}$. A Gaussian blur with standard deviation 1 was applied to the binary shape (Fig. 2D) so that the mean of $\{g_s\}$ $(-0.31)$ is of a comparable to the mean of $\{g_r\}$ $(-0.28)$. Poisson noise was subsequently applied to the blurred mask such that the standard deviation of $\{g_s\}$ (0.10) was comparable to the standard deviation of $\{g_t\}$ (0.09). Scaling the blurred image intensities up by 500 and applying Poisson noise achieved this effect. The resulting noisy image stack was normalized to $[0, 1]$. The plane $z = 0$ was taken as the foreground seed set (the bottom of the shape in Fig. 2A), while background seeds were taken to be the plane $z = 80$ (above the shape in Fig. 2A).

### *C. Cell Tracking Challenge images*

We tested our method on Cell Tracking Challenge datasets Fluo-C3DH-A549 and Fluo-C3DH-A549-SIM (www.celltrackingchallenge.net) [2], [3]. These datasets contain real and simulated 3D confocal images of lung cancer cells, respectively. The simulated images were generated with the FiloGen model-based cell generator [4], [5]. Both datasets have planar pixel width of 0.27 $\mu$m and slice separation 2.3 $\mu$m. Test and training datasets are publicly available from the challenge website, but ground truth segmentations are only available for the training datasets. Our results for the test datasets were externally validated by the Cell Tracking Challenge, which allows a comparison of our results with other state of the art methods in Section IV-B2. The measure used by the Cell Tracking Challenge is based on the Jaccard score, but includes a factor to measure cell detection [3]. The methods compared below all detected the cells perfectly, and therefore this measure is equivalent to the Jaccard score. Additionally, we use our own comparison measures on the training dataset of Fluo-C3DH-A549-SIM to compare our results with the methods presented in Section III-C.

Pre- and post-processing steps were required for the PW, RW, and CERW, and are detailed in the supplementary material. For all three methods, the pre-processing steps were rescaling in $z$ to obtain isotropic resolution, applying a band pass filter, and applying 2D contrast-limited adaptive histogram equalization [6] to the second time series in Fluo-C3DH-A549-SIM. In order to better capture the long and branching filopodia in the second time series of both datasets, a high value of $\kappa$ was used. Because the curvature term is independent of the original image, this led to the filopodia being detected with a wider cross section than desired. Additionally, a higher curvature weighting led to background leaking into the cell through areas of low membrane intensity in some images. Accordingly, the following postprocessing steps are required (for specific parameter values see Supplementary Table III).

The first step was to apply morphological dilation (radius $r_D^{PP}$), fill, and erosion (radius $r_E^{PP}$) to fill any holes inside the cell. The second step was to move the surface of each binary mask to the nearest edge of the original input image. A Gaussian filter with standard deviation $\sigma_p$ was applied to the input image followed by a Sobel filter to obtain the gradient magnitude. A surface mesh was generated from the binary segmentation mask using Matlab's isosurface function. For each vertex $v$ of the mesh, lines of length 10 were drawn along the surface normal from $v$ extending to the interior and exterior of the surface. To prevent intersection with another part of the surface, each line was truncated so that the distance to the surface only increases along the line away from $v$. The voxel $u$ along both of these lines with the highest gradient magnitude was selected as the new location of the surface vertex. If $u$ was interior to the surface, then all voxels intersecting the line between $v$ and $u$ were assigned the background value, while the opposite assignment was made if $u$ was exterior. The volume generated in this manner was smoothed using a median filter of radius 1.

### II. Supplementary tables

#### A. Parameter tables

Parameter values for selecting seeds (Supplementary Table I), band pass segmentation (Supplementary Table II), and post processing of cell tracking challenge data (Supplementary Table III).

Seed selection was not performed on the synthetic test image, but instead seeds were chosen at the top and bottom slices of the image stack. Foreground seeds for real microscopy data were only selected using the maximal surfaces approach, and therefore no dilation or erosion was required for foreground seeds. The foreground seeds for movie 01 of the Cell Tracking Challenge datasets were only generated using dilation and erosion, whereas the foreground seeds for movie 02 of the Cell Tracking Challenge datasets were generated using both the maximal surfaces method and dilation and erosion. All background seeds were selected using dilation and erosion.

Parameter values for seed selection were set manually so that foreground seeds were placed inside the cell being segmented and background seeds placed outside, while aiming to put seeds close to the cell boundary on either side. Many factors play a role in restricting the range of values these parameters can take, including image resolution, fluorescence levels, and cell shape. It was therefore necessary to adjust these values for each dataset, even in cases where the resolution is the same (Cell Tracking Challenge movies).

The band pass segmentation was optimized for the synthetic test image and the real microscopy data using coordinate ascent. The Cell Tracking Challenge training dataset were processed using the software and parameter values found on the Cell Tracking Challenge website (www.celltrackingchallenge.net).

Postprocessing of segmentations was required for the second movie in each of the Cell Tracking Challenge datasets because of the large value of $\kappa$ required to segment the branching filopodia.

SUPPLEMENTARY TABLE I
Parameter Values for Seed Selection

| Parameter[a] | Microscopy data | Cell Tracking Challenge | |
|---|---|---|---|
| | | movie 1 | movie 2[b] |
| Mean filter radius[c] | 2 | 0 | 0(4) |
| Median filter radius[c] | 0 | 2 | 2(4) |
| $w_{\text{phan}}$ | 21 | 21 | 21 |
| $r^d_{BG}$ | 20 | 30 | 30 |
| $r^e_{BG}$ | 5 | 5 | 1 |
| $r^d_{FG}$ | N/A | 20 | 20 |
| $r^e_{FG}$ | N/A | 35 | 40 |

[a] All size measurements are in image voxels
[b] mean and median filters were adjusted depending on the method; the first number represents the size for the dilation-fill-erosion method, while the bracketed number represents the size for the maximal surfaces method.
[c] Filter radius of size 0 indicates no filtering occurred.

SUPPLEMENTARY TABLE II
Parameter Values for Band Pass Segmentation

| Parameter[a] | Microscopy data | Synthetic image |
|---|---|---|
| $w_{\max}$ | 0.0527 | 1.0 |
| $\sigma_B$ | 36.9457 | 84.6729 |
| $\sigma_S$ | 4.0028 | 0.0210 |
| $\alpha$ | 1.1033 | 1.0 |
| $\tau$ | 0.0125 | 0.0011 |

[a] All size measurements are in image voxels

SUPPLEMENTARY TABLE III
Parameter Values for Cell Tracking Challenge Post Processing

| Parameter[a] | Fluo-C3DH-A549 (Movie 02) | Fluo-C3DH-A549-SIM (Movie 02) |
|---|---|---|
| $r_D^{PP}$ | 4 | 5 |
| $r_D^{PP}$ | 4 | 5 |
| $\sigma_p$ | 2.0 | 2.0 |
| $r_{max}$ | 10 | 10 |

[a] All size measurements are in image voxels

### B. Full Cell Tracking Challenge results

Supplementary Table IV is an extension of Table III, with the addition of results from the random forest pixel classifier (RF), power watershed (PW), and band pass (BP).

SUPPLEMENTARY TABLE IV
COMPARISON OF SEGMENTATION METHODS APPLIED TO CELL TRACKING CHALLENGE DATA

| Method[a] | Jaccard | Local Jaccard[b] | | | Hausdorff | Mean BDE[c] | |
|---|---|---|---|---|---|---|---|
| | | $H^-$ | $H^0$ | $H^+$ | | S→GT | GT→S |
| Movie 01 | | | | | | | |
| CNN | 0.875 | 0.734 | 0.788 | 0.674 | **10.8** | 1.34 | 1.22 |
| RF ($N = 50$) | 0.622 | 0.227 | 0.309 | 0.321 | 17.4 | 5.70 | 4.52 |
| RF ($N = 500$) | 0.726 | 0.413 | 0.501 | 0.474 | 16.1 | 3.60 | 2.78 |
| PW ($q = 2$) | 0.877 | 0.773 | 0.795 | 0.673 | 16.6 | 1.64 | 1.15 |
| BP | 0.879 | 0.756 | 0.789 | 0.640 | 20.8 | 1.25 | 1.81 |
| RW ($\beta = 10000$) | 0.899 | **0.811** | 0.826 | 0.694 | 16.4 | **1.02** | 1.24 |
| CERW ($\beta = 2000, \kappa = 0.80$) | **0.902** | 0.805 | **0.832** | **0.713** | 13.4 | 1.03 | **1.09** |
| CERW ($\beta = 9000, \kappa = 0.40$) | 0.881 | 0.765 | 0.799 | 0.687 | *13.0* | 1.42 | 1.14 |
| Movie 02 | | | | | | | |
| CNN | **0.821** | **0.738** | **0.736** | **0.666** | **16.5** | **1.47** | **1.14** |
| RF ($N = 50$) | 0.465 | 0.218 | 0.303 | 0.334 | 20.4 | 5.89 | 3.99 |
| RF ($N = 500$) | 0.623 | 0.392 | 0.474 | 0.475 | *18.1* | 4.02 | 2.76 |
| PW ($q = 2$) | 0.692 | 0.541 | 0.499 | 0.407 | 67.4 | 3.02 | 6.07 |
| BP | 0.751 | 0.635 | 0.591 | 0.495 | 41.1 | 2.00 | 3.52 |
| RW ($\beta = 8000$) | 0.708 | 0.520 | 0.508 | 0.431 | 65.1 | 2.71 | 6.29 |
| CERW ($\beta = 5500, \kappa = 0.40$) | 0.643 | 0.438 | 0.503 | 0.479 | 27.9 | 3.75 | 3.51 |
| CERW post-processed[d]($\beta = 5500, \kappa = 0.40$) | *0.788* | *0.671* | *0.643* | *0.572* | 27.4 | *1.84* | *1.91* |

[a] CNN: pretrained convolutional neural network RF: random forest classifier; BP: band pass method; PW: power watershed; RW: random walker; CERW: curvature-enhanced random walker.
[b] $H^-$: highly negative curvature; $H^0$: low curvature; $H^+$: highly positive curvature.
[c] Boundary displacement error. S→GT: mean distance from segmentation to ground truth; GT→S: mean distance from ground truth to segmentation.
[d] Post processing to improve precision is described in Section III-D2.
Bold numbers highlight the best value, italic numbers highlight the best non-cnn value.

### C. Inter-annotator variability

The inter-annotator variability (Supplementary Tables V–VIII) was generated for comparison with results from the segmentation methods (Table IV, Supplementary Tables IX–XI). The largest difference between annotators was taken to be a limit on segmentation performance comparison, such that any segmentations showing a lower difference than this limit is deemed optimal, and any differences in scores beyond this bound cannot be taken to be real differences in the segmentation performance in comparison to annotator 1. Comparing the results Table IV with the corresponding inter-annotator variability (Supplementary Table V), the random forest classifier, band pass method, and power watershed all score worse than the bounding values of all measures, suggesting that the curvature-enhanced random walker does show a real improvement on these methods across all measures. The random walker only scores worse than the bounding values in the Hausdorff distance and local Jaccard score for areas of highly positive curvature. This suggests that the curvature-enhanced random walker performs better than the random walker in areas of highly positive curvature and in Hausdorff distance, but doesn't offer any real improvement in other areas in comparison to annotator 1. In order to determine if alternative interpretations yield different results, it is necessary to compare the segmentation results to alternative manual segmentations.

SUPPLEMENTARY TABLE V
MANUAL SEGMENTATION COMPARISON WITH ANNOTATOR 1

| Method[a] | Jaccard | Local Jaccard[b] | | | Hausdorff | Mean BDE[c] | |
|---|---|---|---|---|---|---|---|
| | | $H^-$ | $H^0$ | $H^+$ | | S→GT | GT→S |
| Annotator 2 | 0.817 | 0.744 | 0.784 | 0.740 | 7.33 | 1.14 | 1.20 |
| Annotator 3 | 0.891 | 0.847 | 0.862 | 0.814 | 6.43 | 0.544 | 0.687 |
| Aggregated | 0.945 | 0.835 | 0.890 | 0.855 | 5.19 | 0.300 | 0.355 |

[a] Aggregated segmentation generated using majority voting.
[b] $H^-$: highly negative curvature; $H^0$: low curvature; $H^+$: highly positive curvature.
[c] Boundary displacement error. S→GT: mean distance from segmentation to ground truth; GT→S: mean distance from ground truth to segmentation.

SUPPLEMENTARY TABLE VI

MANUAL SEGMENTATION COMPARISON WITH ANNOTATOR 2

| Method[a] | Jaccard | Local Jaccard[b] | | | Hausdorff | Mean BDE[c] | |
|---|---|---|---|---|---|---|---|
| | | $H^-$ | $H^0$ | $H^+$ | | S→GT | GT→S |
| Annotator 1 | 0.817 | 0.751 | 0.777 | 0.690 | 7.33 | 1.19 | 1.14 |
| Annotator 3 | 0.817 | 0.766 | 0.776 | 0.707 | 8.38 | 0.991 | 1.58 |
| Aggregated | 0.866 | 0.830 | 0.839 | 0.794 | 6.75 | 0.758 | 0.785 |

[a] Aggregated segmentation generated using majority voting.
[b] $H^-$: highly negative curvature; $H^0$: low curvature; $H^+$: highly positive curvature.
[c] Boundary displacement error. S→GT: mean distance from segmentation to ground truth; GT→S: mean distance from ground truth to segmentation.

SUPPLEMENTARY TABLE VII

MANUAL SEGMENTATION COMPARISON WITH ANNOTATOR 3

| Method[a] | Jaccard | Local Jaccard[b] | | | Hausdorff | Mean BDE[c] | |
|---|---|---|---|---|---|---|---|
| | | $H^-$ | $H^0$ | $H^+$ | | S→GT | GT→S |
| Annotator 1 | 0.891 | 0.843 | 0.867 | 0.795 | 6.43 | 0.687 | 0.544 |
| Annotator 2 | 0.817 | 0.760 | 0.792 | 0.748 | 8.38 | 1.16 | 0.991 |
| Aggregated | 0.943 | 0.941 | 0.932 | 0.905 | 4.41 | 0.344 | 0.238 |

[a] Aggregated segmentation generated using majority voting.
[b] $H^-$: highly negative curvature; $H^0$: low curvature; $H^+$: highly positive curvature.
[c] Boundary displacement error. S→GT: mean distance from segmentation to ground truth; GT→S: mean distance from ground truth to segmentation.

SUPPLEMENTARY TABLE VIII

MANUAL SEGMENTATION COMPARISON WITH AGGREGATED ANNOTATION

| Method[a] | Jaccard | Local Jaccard[b] | | | Hausdorff | Mean BDE[c] | |
|---|---|---|---|---|---|---|---|
| | | $H^-$ | $H^0$ | $H^+$ | | S→GT | GT→S |
| Annotator 1 | 0.945 | 0.858 | 0.888 | 0.847 | 5.19 | 0.355 | 0.300 |
| Annotator 2 | 0.866 | 0.820 | 0.848 | 0.816 | 6.75 | 0.785 | 0.758 |
| Annotator 3 | 0.943 | 0.927 | 0.929 | 0.893 | 4.41 | 0.238 | 0.344 |

[a] Aggregated segmentation generated using majority voting.
[b] $H^-$: highly negative curvature; $H^0$: low curvature; $H^+$: highly positive curvature.
[c] Boundary displacement error. S→GT: mean distance from segmentation to ground truth; GT→S: mean distance from ground truth to segmentation.

#### D. *Comparison of manual segmentations with automated segmentation methods*

Supplementary Tables IX–XI show the comparison of the segmentation results with each of the two additional annotators, and with the aggregated result. These tables show similar results to that of Table IV, with the exception that here the Hausdorff distance is lower for the random forest classifier than our method. The reduced Hausdorff distance in the random forest classifier is likely due to oversegmentation.

Importantly, these tables show that the curvature-enhanced random walker outperforms the random walker in all measures against all annotators, suggesting that, in all interpretations, the curvature-enhanced random walker provides a more accurate segmentation than the random walker. This observation provides a stronger result than can be seen when comparing to a single annotator, where segmentation performance is tied to a specific interpretation, as seen in Section II-C.

SUPPLEMENTARY TABLE IX
COMPARISON OF SEGMENTATION METHODS WITH ANNOTATOR 2.

| Method[a] | Jaccard | Local Jaccard[b] | | | Hausdorff | Mean BDE[c] | |
|---|---|---|---|---|---|---|---|
| | | $H^-$ | $H^0$ | $H^+$ | | S→GT | GT→S |
| CNN | 0.822 | **0.769** | **0.802** | 0.740 | 11.5 | 1.22 | **0.96** |
| RF ($N = 50$) | 0.730 | 0.647 | 0.696 | 0.663 | 10.2 | 2.16 | 1.95 |
| RF ($N = 500$) | 0.740 | 0.661 | 0.711 | 0.675 | **10.1** | 2.05 | 1.79 |
| PW ($q = 2$) | 0.800 | 0.740 | 0.751 | 0.676 | 14.4 | 1.37 | 1.117 |
| BP | 0.731 | 0.636 | 0.682 | 0.634 | 10.9 | 2.01 | 2.158 |
| RW ($\beta = 4900$) | 0.827 | 0.759 | 0.787 | 0.719 | 15.9 | 1.06 | 1.14 |
| CERW ($\beta = 4900, \kappa = 0.06$) | **0.833** | 0.764 | 0.799 | 0.754 | 11.9 | 1.04 | 1.00 |
| CERW ($\beta = 3500, \kappa = 0.12$) | **0.833** | 0.764 | 0.800 | **0.756** | 10.6 | **1.03** | 1.00 |
| CERW ($\beta = 2500, \kappa = 0.18$) | 0.832 | 0.762 | 0.799 | 0.755 | 10.7 | 1.03 | 1.02 |
| CERW ($\beta = 1500, \kappa = 0.30$) | 0.830 | 0.757 | 0.796 | 0.751 | 11.0 | 1.05 | 1.07 |
| CERW $(1000, 0.50)$ | 0.826 | 0.751 | 0.794 | 0.751 | 11.4 | 1.08 | 1.101 |

[a] CNN: pretrained convolutional neural network; RF: random forest classifier; PW: power watershed; BP: band pass method; RW: random walker; CERW: curvature-enhanced random walker.
[b] $H^-$: highly negative curvature; $H^0$: low curvature; $H^+$: highly positive curvature.
[c] Boundary displacement error. S→GT: mean distance from segmentation to ground truth; GT→S: mean distance from ground truth to segmentation.
Bold numbers highlight the best value.

SUPPLEMENTARY TABLE X
COMPARISON OF SEGMENTATION METHODS WITH ANNOTATOR 3

| Method[a] | Jaccard | Local Jaccard[b] | | | Hausdorff | Mean BDE[c] | |
|---|---|---|---|---|---|---|---|
| | | $H^-$ | $H^0$ | $H^+$ | | S→GT | GT→S |
| CNN | 0.864 | 0.831 | 0.837 | 0.772 | **7.52** | **0.708** | 0.734 |
| RF ($N = 50$) | 0.811 | 0.707 | 0.763 | 0.729 | 8.01 | 1.33 | 1.27 |
| RF ($N = 500$) | 0.820 | 0.727 | 0.779 | 0.751 | 8.28 | 1.19 | 1.20 |
| PW ($q = 2$) | 0.794 | 0.750 | 0.735 | 0.640 | 11.8 | 1.26 | 1.157 |
| BP | 0.809 | 0.731 | 0.753 | 0.667 | 13.2 | 1.59 | 1.10 |
| RW ($\beta = 4900$) | 0.848 | 0.812 | 0.801 | 0.687 | 19.4 | 1.168 | 0.711 |
| CERW ($\beta = 4900, \kappa = 0.06$) | 0.875 | 0.839 | 0.841 | 0.763 | 11.5 | 0.779 | 0.586 |
| CERW ($\beta = 3500, \kappa = 0.12$) | **0.879** | **0.843** | **0.847** | **0.775** | 10.9 | 0.748 | **0.566** |
| CERW ($\beta = 2500, \kappa = 0.18$) | 0.876 | 0.838 | 0.842 | 0.766 | 11.4 | 0.789 | 0.578 |
| CERW ($\beta = 1500, \kappa = 0.30$) | 0.870 | 0.831 | 0.834 | 0.754 | 12.2 | 0.857 | 0.603 |
| CERW $(1000, 0.50)$ | 0.874 | 0.833 | 0.839 | 0.763 | 12.7 | 0.836 | 0.593 |

[a] CNN: pretrained convolutional neural network; RF: random forest classifier; PW: power watershed; BP: band pass method; RW: random walker; CERW: curvature-enhanced random walker.
[b] $H^-$: highly negative curvature; $H^0$: low curvature; $H^+$: highly positive curvature.
[c] Boundary displacement error. S→GT: mean distance from segmentation to ground truth; GT→S: mean distance from ground truth to segmentation.
Bold numbers highlight the best value.

SUPPLEMENTARY TABLE XI
COMPARISON OF SEGMENTATION METHODS WITH AGGREGATED MANUAL SEGMENTATION (MAJORITY VOTING)

| Method[a] | Jaccard | Local Jaccard[b] | | | Hausdorff | Mean BDE[c] | |
|---|---|---|---|---|---|---|---|
| | | $H^-$ | $H^0$ | $H^+$ | | S→GT | GT→S |
| CNN | 0.878 | 0.848 | 0.860 | 0.777 | 9.28 | 0.743 | 0.626 |
| RF ($N = 50$) | 0.810 | 0.694 | 0.767 | 0.717 | 8.21 | 1.30 | 1.36 |
| RF ($N = 500$) | 0.825 | 0.676 | 0.764 | 0.694 | **7.78** | 1.25 | 1.16 |
| PW ($q = 2$) | 0.812 | 0.778 | 0.762 | 0.648 | 13.4 | 1.11 | 1.20 |
| BP | 0.811 | 0.713 | 0.761 | 0.662 | 11.4 | 1.52 | 1.22 |
| RW ($\beta = 4900$) | 0.868 | 0.838 | 0.830 | 0.701 | 17.7 | 0.944 | 0.705 |
| CERW ($\beta = 4900, \kappa = 0.06$) | 0.894 | 0.860 | 0.866 | 0.792 | 10.2 | 0.639 | 0.582 |
| CERW ($\beta = 3500, \kappa = 0.12$) | **0.896** | **0.862** | **0.871** | **0.803** | 9.02 | **0.615** | **0.564** |
| CERW ($\beta = 2500, \kappa = 0.18$) | 0.894 | 0.858 | 0.867 | 0.795 | 10.7 | 0.647 | 0.574 |
| CERW ($\beta = 1500, \kappa = 0.30$) | 0.888 | 0.849 | 0.859 | 0.780 | 11.6 | 0.708 | 0.599 |
| CERW $(1000, 0.50)$ | 0.889 | 0.845 | 0.861 | 0.787 | 11.4 | 0.698 | 0.604 |

[a] CNN: pretrained convolutional neural network; RF: random forest classifier; PW: power watershed; BP: band pass method; RW: random walker; CERW: curvature-enhanced random walker.
[b] $H^-$: highly negative curvature; $H^0$: low curvature; $H^+$: highly positive curvature.
[c] Boundary displacement error. S→GT: mean distance from segmentation to ground truth; GT→S: mean distance from ground truth to segmentation.
Bold numbers highlight the best value.

### *E. Processing times*

A comparison of the average time required to segment image volumes from each dataset with each method is provided in Table XII. The CERW is the slowest method in all cases, with the larger and more complex image volumes in the second movie of the Cell Tracking Challenge data requiring roughly 20 times longer to process than the slowest non-random walker-based method (the PW). However, roughly half of the computation time for the CERW is taken up by the initial RW segmentation. We simulated diffusion for this step, in order to provide continuity between methods, but faster methods for computing the initial RW values are available [7], [8], which would greatly improve the speed. Furthermore, the method of simulation for the curvature enhancement step has not been optimized for time-efficiency, and could potentially be improved upon to reduce the segmentation time. Finally, the seed selection also requires a significant proportion of the processing time. However, this step was not performed on a GPU, which could significantly reduce computation time. Note that because the pretrained neural network could not be implemented on the NVIDIA Tesla-K80 GPU, the time required to segment the images using this method took significantly longer than would be expected for a CNN, and we have therefore omitted these timings.

SUPPLEMENTARY TABLE XII
AVERAGE TIMINGS OF SEGMENTATION METHODS

| Method[a] | Synthetic image | Microscopy data | Cell Tracking Challenge[b] | |
|---|---|---|---|---|
| | $(160 \times 160 \times 80)$[c] | $(120 \times 120 \times 100)$[c] | 01 $(140 \times 150 \times 160)$[c] | 02 $(280 \times 350 \times 300)$[c] |
| RF ($N = 50$) | 24.0 | 51.2 | 36.0 | 44.4 |
| RF ($N = 500$) | 47.0 | 57.7 | 31.5 | 51.8 |
| BP[d] | 4.0 | 10.8 | 1.2 | 1.6 |
| Seeding[e] | N/A | 146.5 | 33.2 | 158.0 |
| PW ($q = 2$) | 12.0 | 19.6 | 121.9 | 144.6 |
| RW[d] | 125.0 | 40.6 | 160.4 | 1600 |
| CERW[d] | 565.0 | 74.8 | 210.5 | 2887 |

[a] RF: random forest classifier; BP: band pass and threshold; PW: power watershed; RW: random walker; CERW: curvature-enhanced random walker.
[b] Dataset Fluo-C3DH-A549-SIM taken from the Cell Tracking Challenge ( [2], [3]).
[c] Dimensions of a typical image volume after cropping.
[d] Timings are for the parameter values yielding the optimal Jaccard score for each dataset.
[e] Time required to generate seeds for graph-based methods
Times represent the average time required to segment one image volume, in seconds.

### III. Supplementary Figures

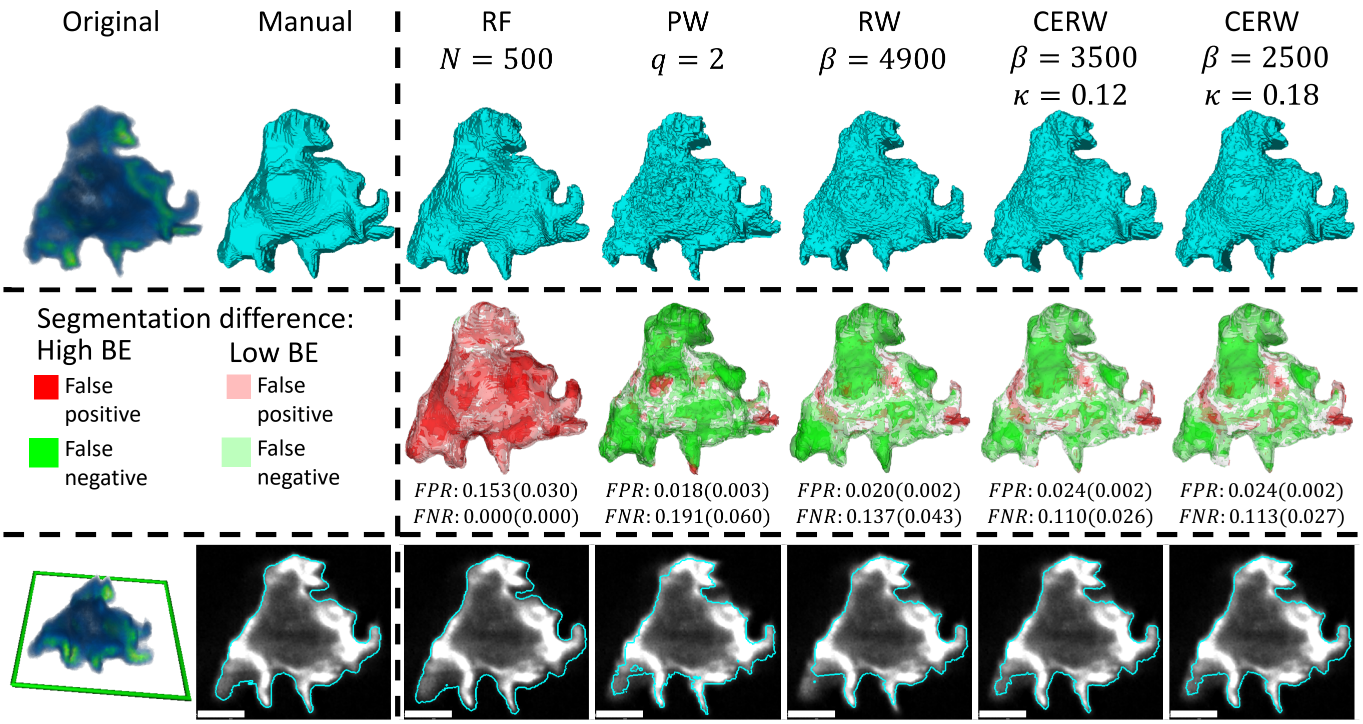


Supplementary Fig. 3. Comparison between manual segmentation and random forest pixel classifier (RF), power watershed (PW), random walker (RW), and curvature-enhanced random walker (CERW). Top row: surfaces of each segmentation. Middle row: differences between segmentation results and manual segmentation, where red marks false positive, green marks false negative, and reduced opacity is given to pixels with a boundary error of 1 (low BE). FPR: false positive rate, FNR: false negative rate, numbers in brackets are error rates of high BE points only. Bottom row: slices through each surface compared to original image, scale bars represent 5 $\mu$m.

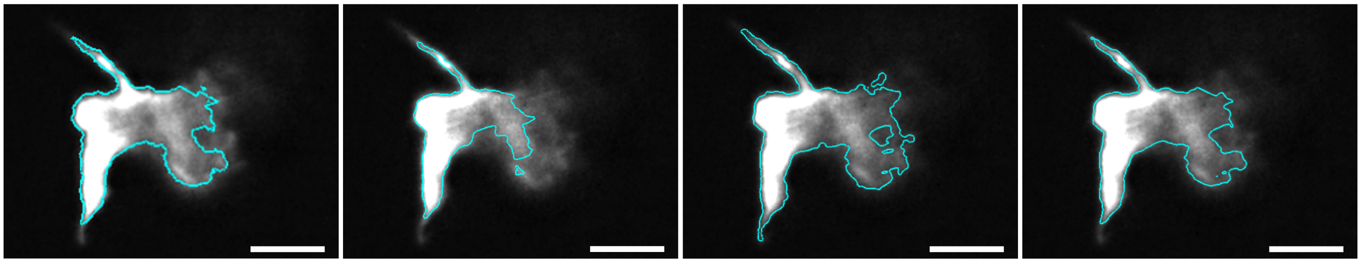

Supplementary Fig. 4. Alternative manual segmentations for the slice shown in Fig 7. From left to right: segmentations from annotator 1, annotator 2, annotator 3, and the aggregated segmentation. The most notable disagreements between annotators are in the length of the protrusions (top and bottom of the image) and the position of the boundary to the right of the image. Annotator 1, which features in Fig. 7 lies between the other two annotators, suggesting that this is a reasonable interpretation of the segmentation boundary.

Supplementary Fig. 5. 1–11: Segmentations of all microscopy images, showing the optimal segmentations with the standard random walker (blue) and the curvature-enhanced random walker (red). Parameter values used to generate these segmentations were $\beta = 4900$ for the standard random walker and $\beta = 3500$, $\kappa = 0.12$ for the curvature-enhanced random walker. These values maximized the mean local Jaccard score for high-curvature areas. Note that the differences in segmentation are most obvious in protrusions (filopodia) present in experiments 1–4, since these structures are finer in the images that were segmented. Grey arrows indicate areas where the curvature enhancement has improved the segmentation.

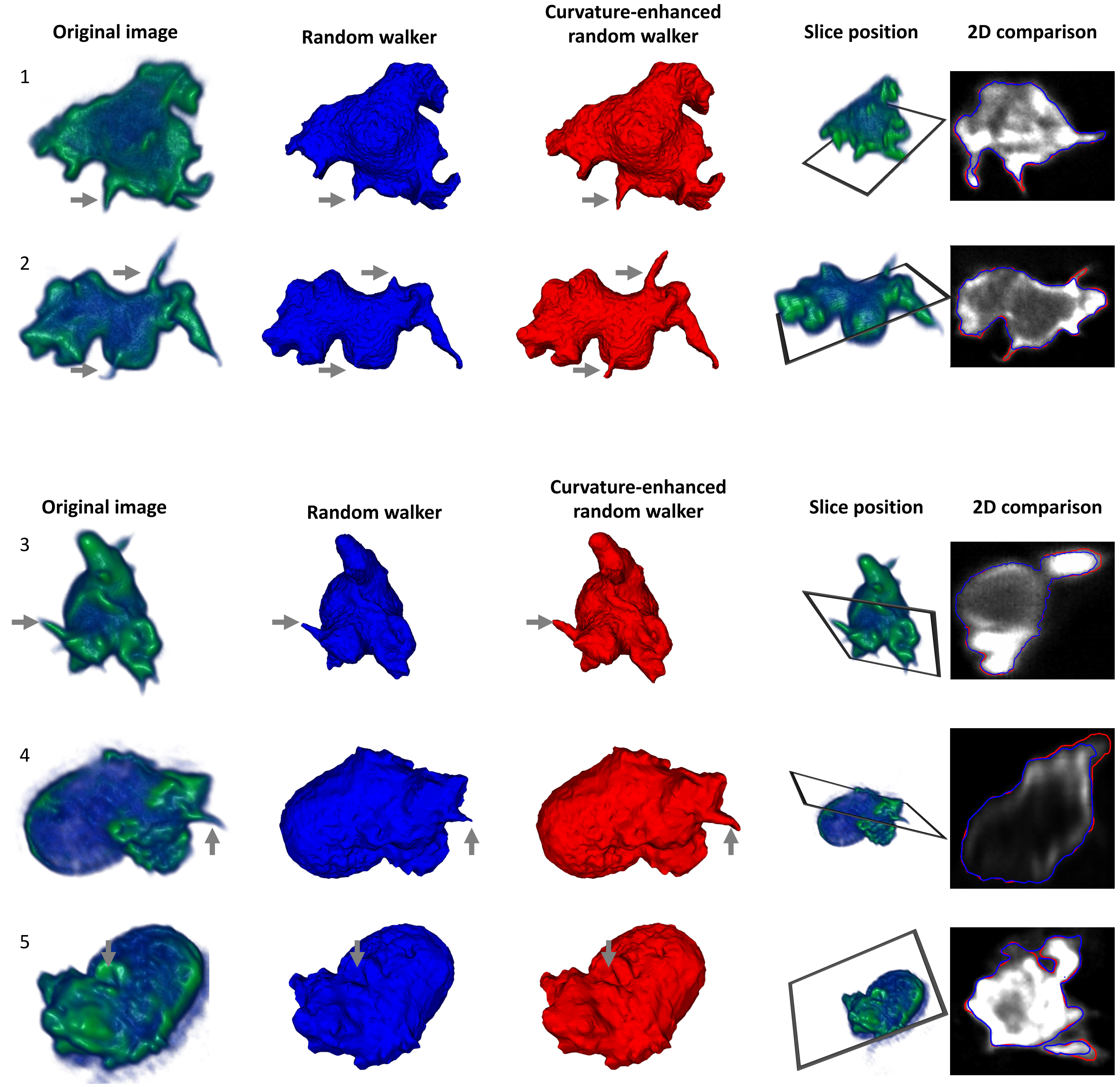

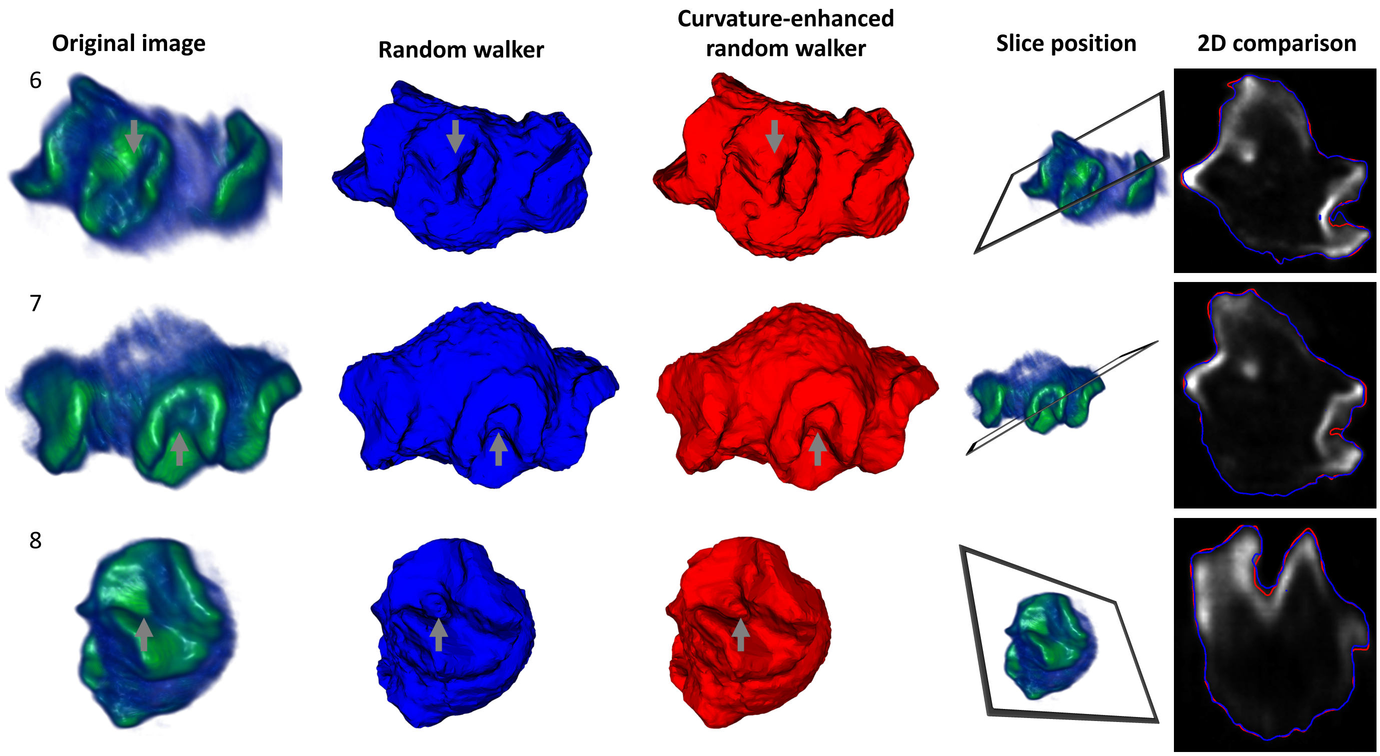
Original image
Random walker
Curvature-enhanced random walker
Slice position
2D comparison
6
7
8

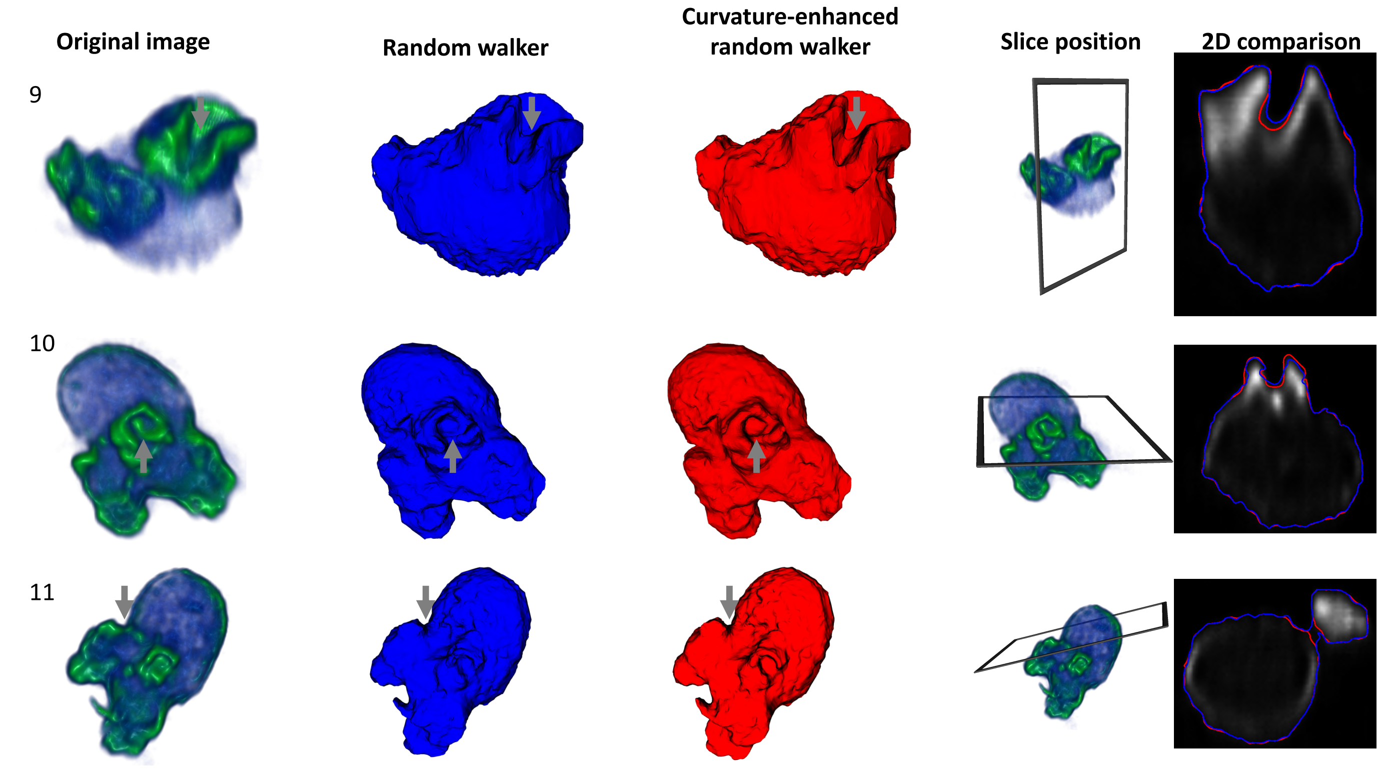
Original image
Random walker
Curvature-enhanced random walker
Slice position
2D comparison
9
10
11
