## Supplementary material for "A Curvature-Enhanced Random Walker Segmentation Method for Detailed Capture of 3D Cell Surface Membranes": Manual Segmentation Protocol

### Manual annotation with Slicer

### Step 1: load data

Download 3D Slicer from www.slicer.org and launch.

1. Click the open icon (a)
2. Click choose File(s) to Add (b)
3. Select files and click OK (c)

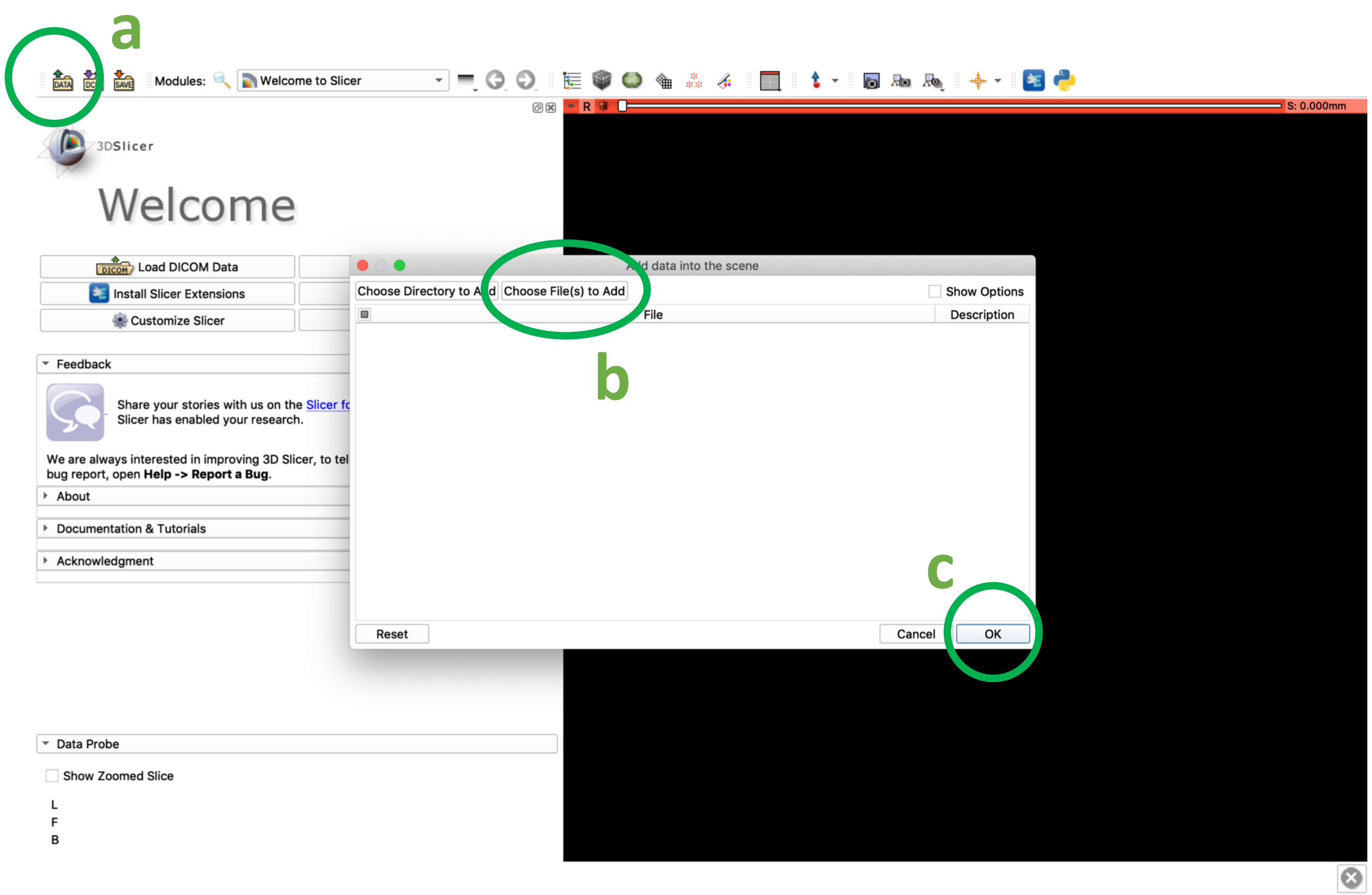

### Step 2: set up segmentation module

1. Select the "Segment Editor" module (a)
2. Click "Add" to create a new segmentation (b)

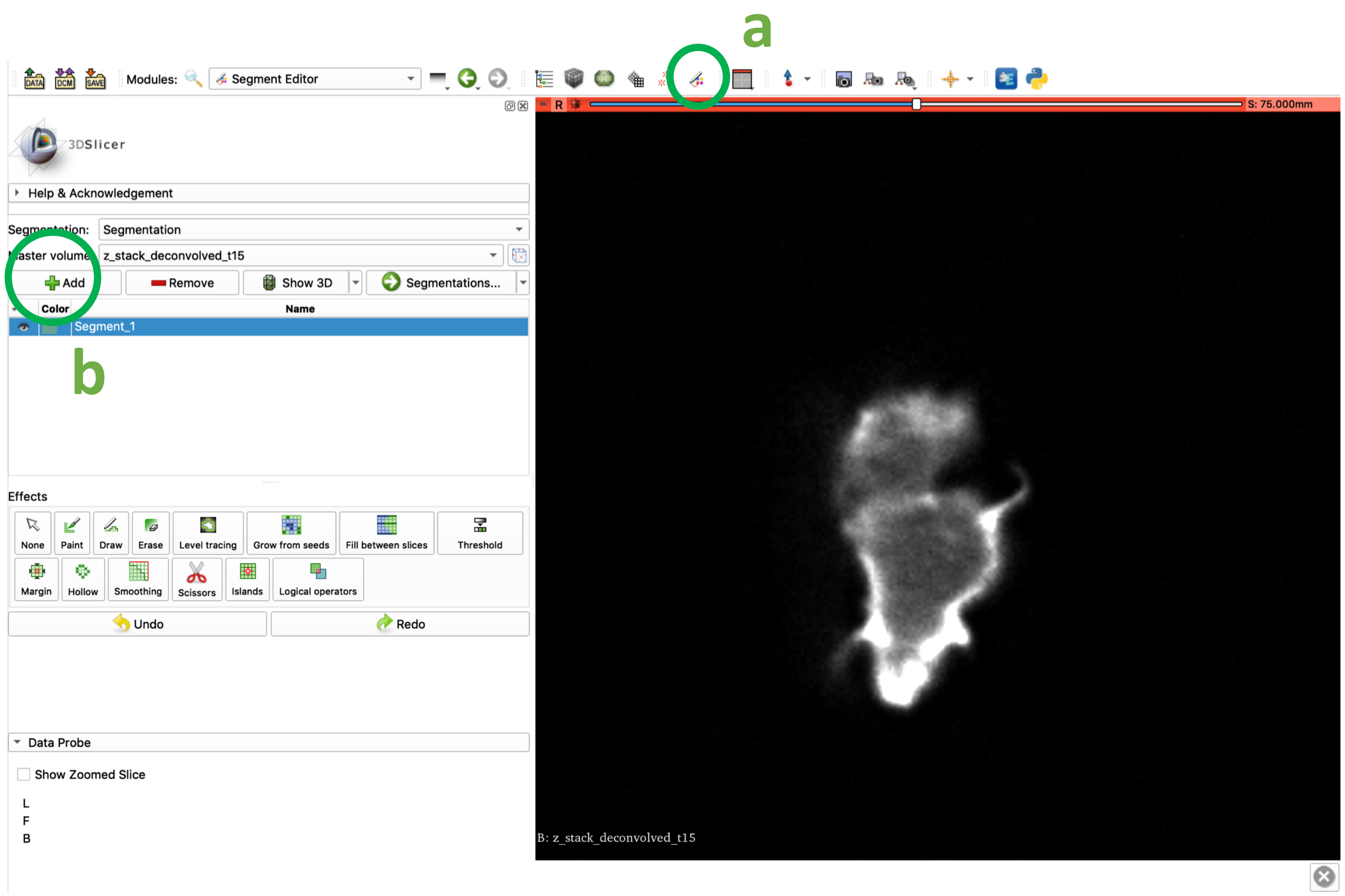

### Step 3: select slice view

- Click the view button (green circle) and select “Red slice only” to display the (x,y)-plane in the viewing pane.

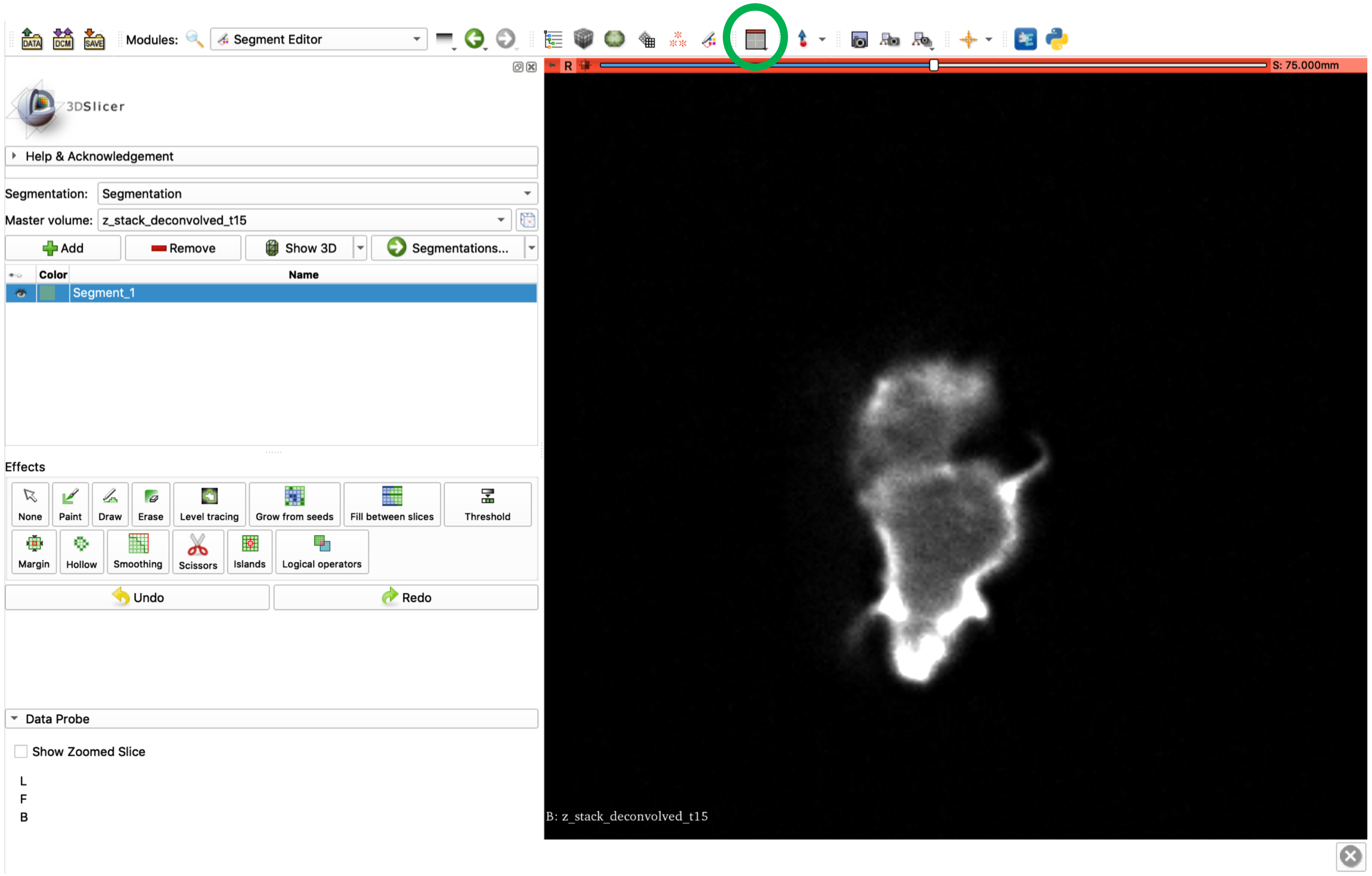

### Step 4: segmentation

- Segment each slice, using left/right or up/down to cycle through the cell.
- The paint tool is most useful for capturing finer details, and brush size can be adjusted either in the Segment Editor panel or by holding shift and scrolling.
- The erase tool functions similarly.
- To zoom in and out, right click and drag with the mouse.
- To move the viewing position (pan), hold shift and left click and drag.

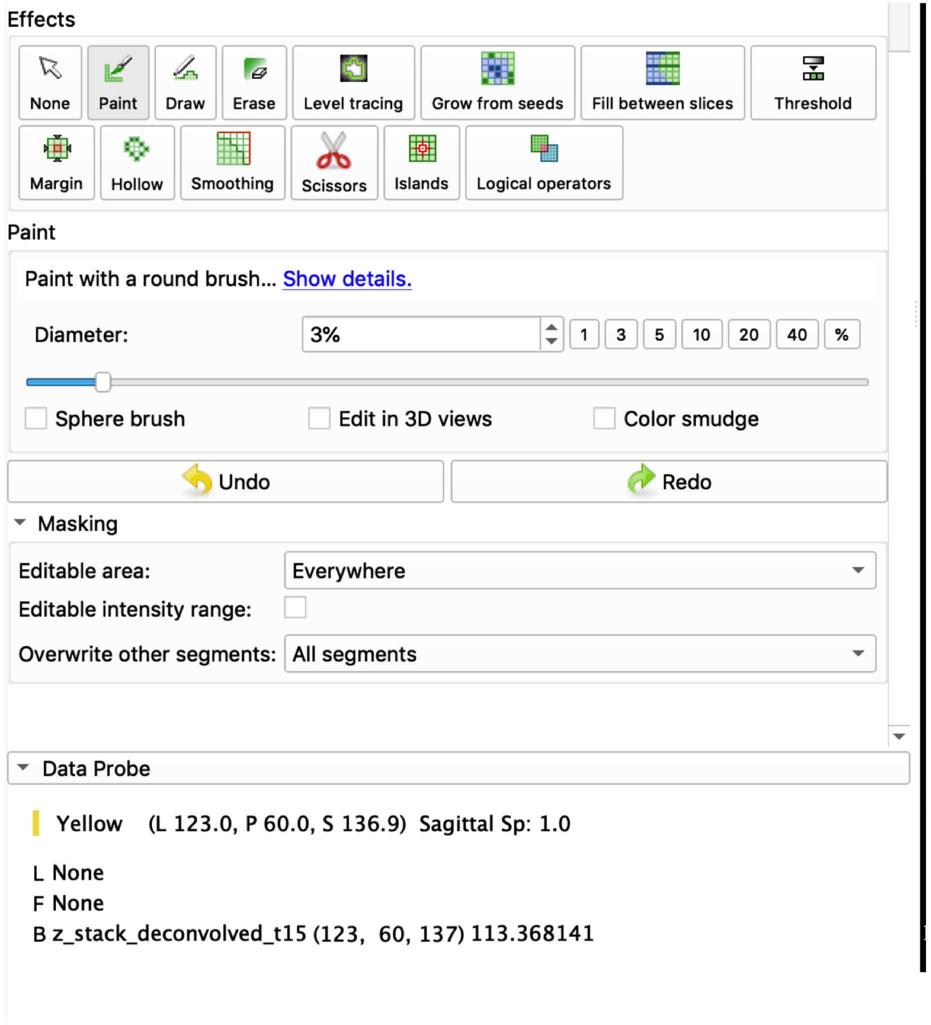

### Step 4: segmentation

- Other tools:
  - Draw: draw an outline contour to fill, can be useful for filling in image, but may not be faster than paint with a large brush.
  - Smoothing: using 3x3x3 median filter before changing views (Step 5) is useful to smooth between slices.
  - Automated methods: not recommended. While they could potentially save time, they may also bias the segmentation choices.

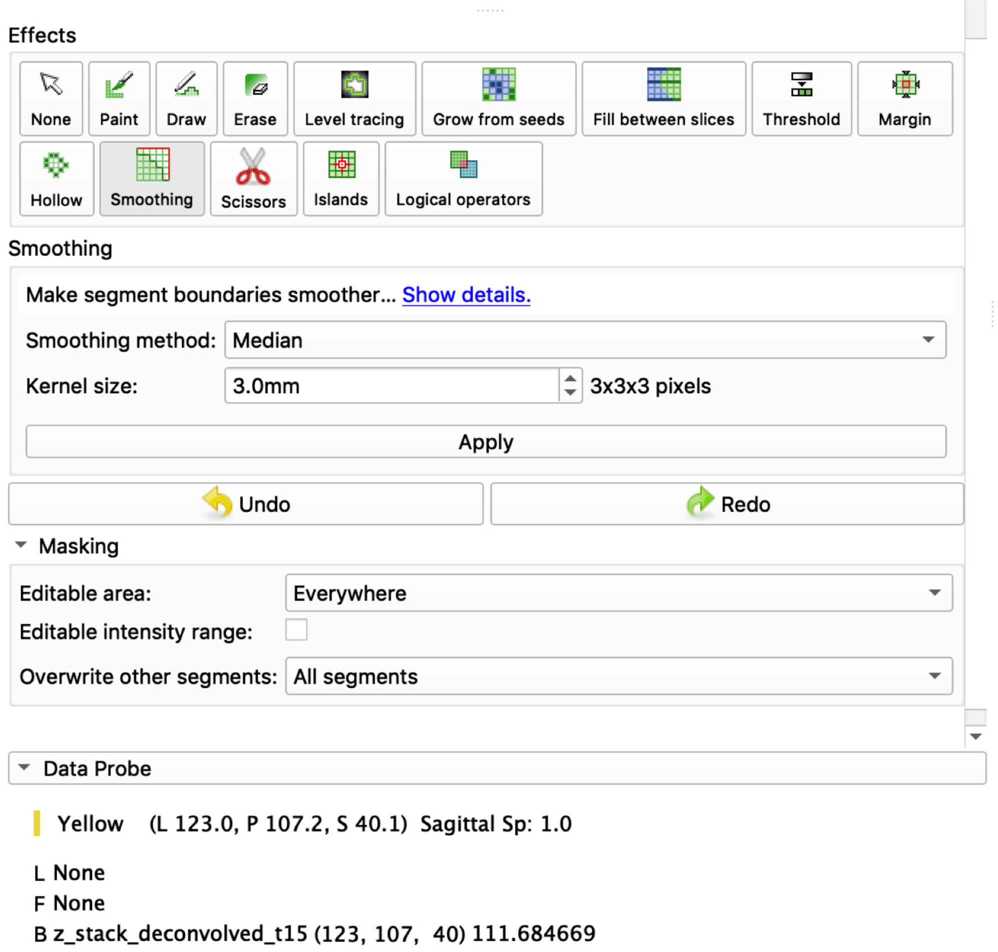

### Step 4: segmentation - tips

- The segmentation appearance can be changed between overlay and outline by clicking on the pin in the top-left corner of the slice pane (a), clicking the arrows below it (b), and clicking the icon next to “Segmentation” (c).
- The colour can be changed by double-clicking the Color square (d), and selecting a new colour using the Color square in the dialog.

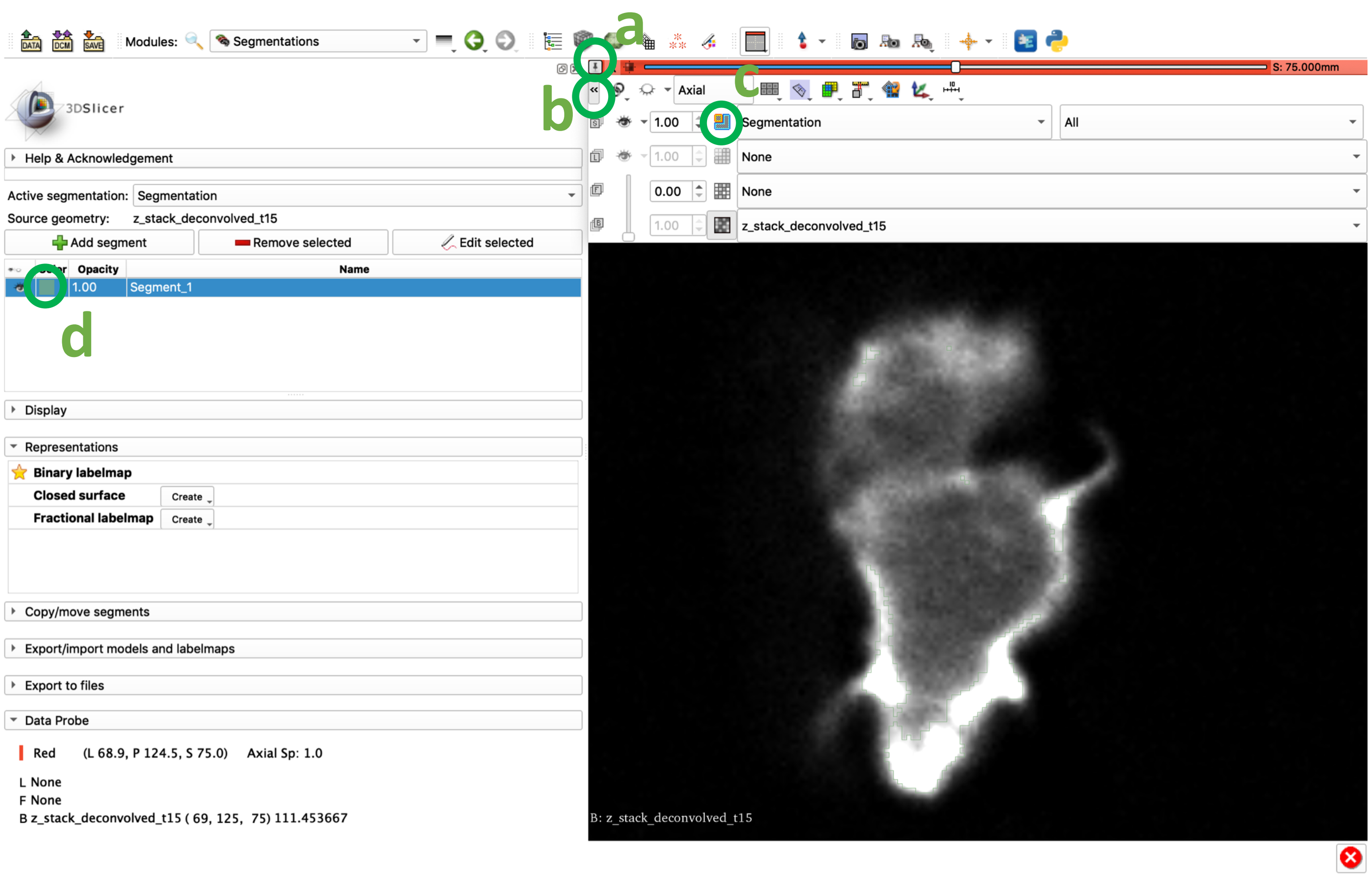

### Step 4: segmentation - tips

- The outline thickness can be adjusted in the "Segmentations" module, which can be accessed from the "Segment Editor" by clicking the "Segmentations…" button.
- To increase the line thickness go to the "Advanced" panel and increase the value in the "Slice intersection thickness" (units are in pixels of the viewer, not pixels of the actual image).

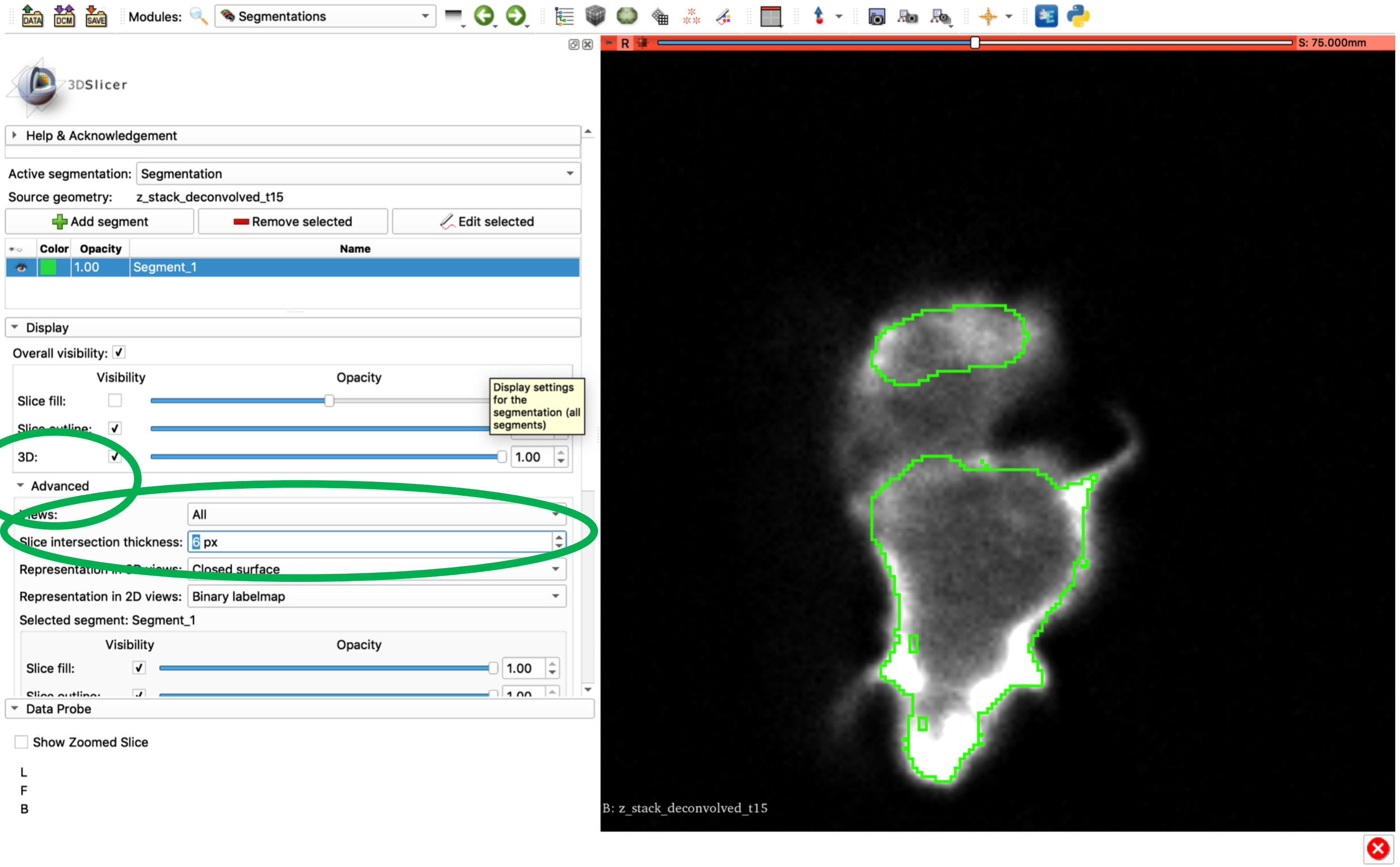

### Step 5: repeat with other plane views

- Click the view button (green circle) and select “Yellow slice only” to display the (y,z)-plane in the viewing pane.
- Repeat step 4, adjusting the previously drawn segmentation where necessary.
- Do the same with the view set to “Green slice only”.

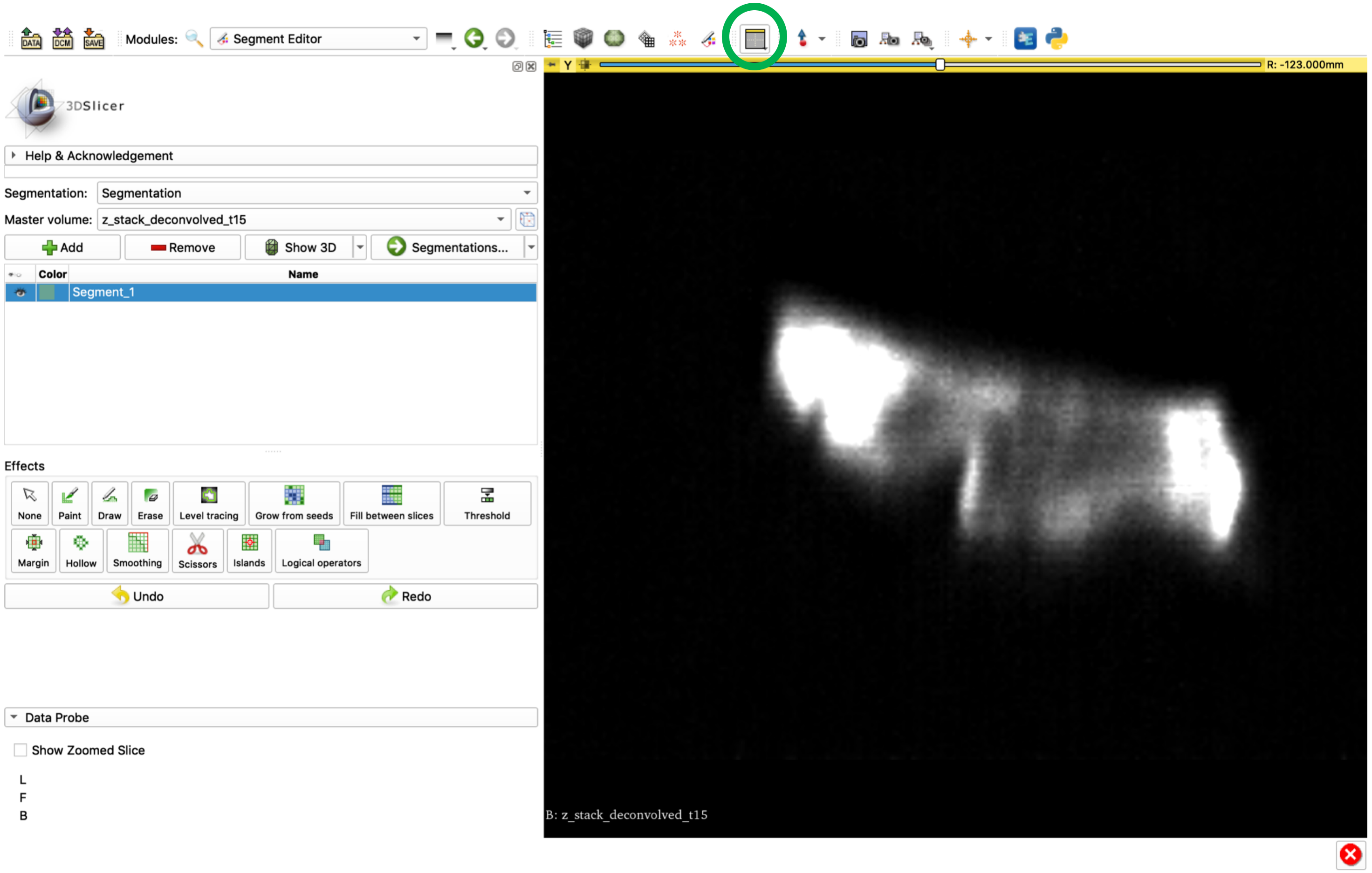

### Step 6: adjust for all slide views

- Show all slice views by selecting “Four up” from the view button (green circle).
- Check segmentation by scrolling through each slice view.
- If the segmentation doesn’t fit in one plane, hold shift and hover the mouse over the area in question to align all other planes to that area.
- Adjust segmentation to best fit all three slice views.

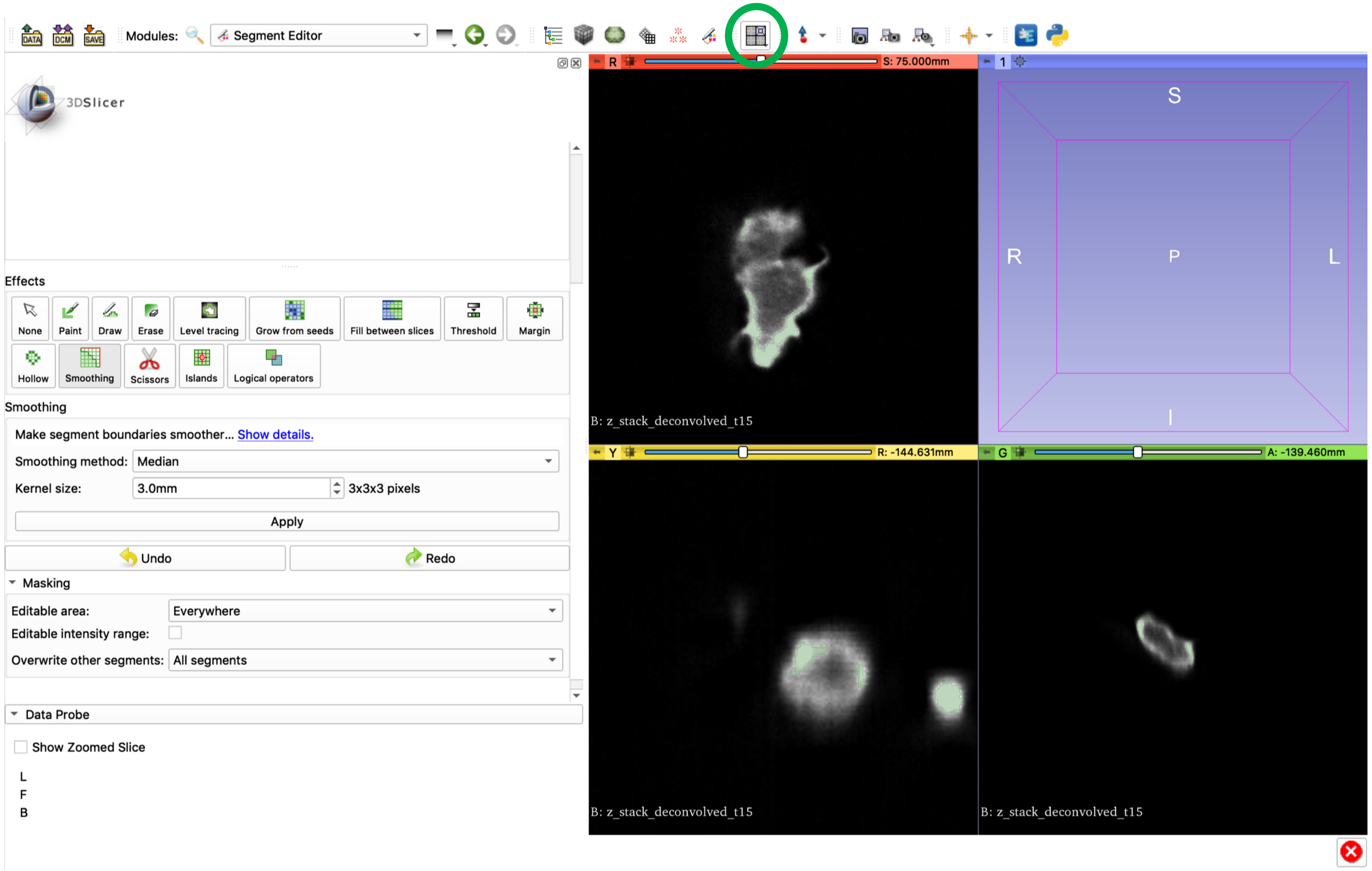

### Step 7: save

- Open the “Segmentations” module (access from “Segment Editor” using the “Segmentations…” button).
- In the “Exoprt/Import models and labelmaps” panel (a), make sure “Export” and “Labelmap” are checked, and click “Export”.
- Click “Save” (b), choose a directory to store the Slicer files and the output, and rename the file “Segmentation-label.nrrd” to “<input_file>_man_seg_<user_initials>.nrrd” (click on the name to edit).
- This output file can be opened in Fiji to confirm it has saved correctly.
- (progress can also be saved through the save menu. To return to the segmentation when reopening Slicer, load the “.mrml” file).

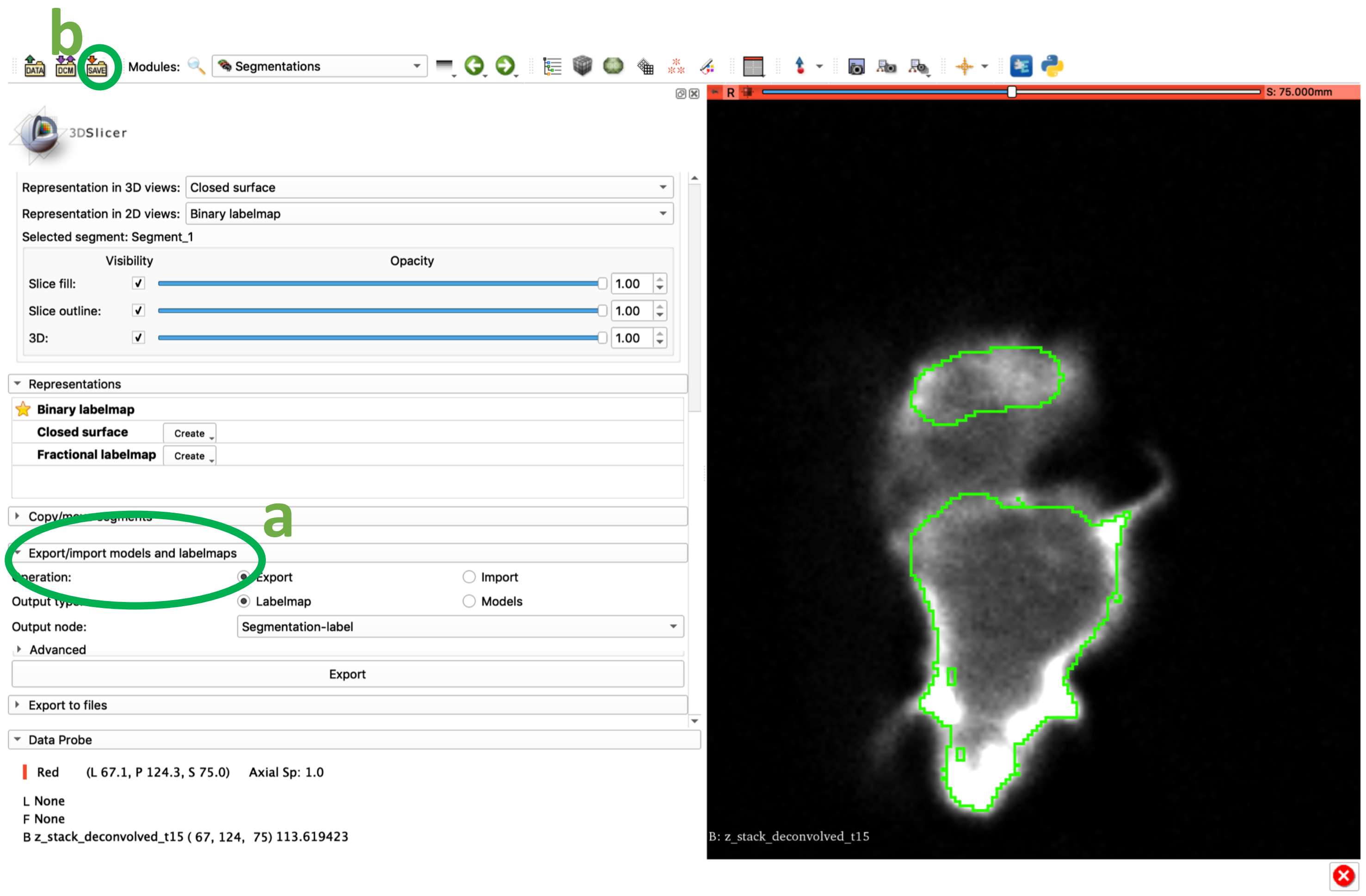

### Additional notes

- Each segmentation should cover the whole cell, but the outline should be as close to the outside of the actin cortex as possible.
- The segmented cells should be fully connected, and should not contain holes.
- Careful attention should be paid to accurately segmenting fine structures, as these are of particular interest in the segmentation.
